# Epigenetic Shifts Reveal Alzheimer’s Origins after Sustained Picomolar Aβ Exposure

**DOI:** 10.1101/2025.08.12.668749

**Authors:** Andrea Paquola, Erica Acquarone, Agnieszka Staniszewski, Andrew F. Teich, Ottavio Arancio

## Abstract

Recent advances in sequencing have identified genetic risk factors for Alzheimer’s disease (AD), but the molecular mechanisms triggering disease onset remain unclear. While high brain levels of amyloid-beta (Aβ) impair synaptic function and memory, exposure to low picomolar (pM) Aβ concentrations - typical of healthy brains - enhances these functions, while sustained exposure results in impairment. To investigate this transition from physiological to pathological Aβ activity, we profiled DNA methylation and gene expression in C57B16 mice subjected to prolonged pM Aβ42 exposure. We identified differentially methylated and expressed genes, including those involved in synaptic function, associated with three phases: memory enhancement (brief exposure), a transitional state with intact memory (intermediate), and memory decline (prolonged). These gene sets may represent early molecular drivers of AD pathogenesis.

## Main text

Amyloid-beta (Aβ) peptide is closely associated with synaptic dysfunction and memory loss in Alzheimer’s disease (AD), with fibrillar Aβ aggregates forming plaques long considered a hallmark of pathology. However, recent evidence highlights the pathogenic role of soluble Aβ species over fibrillary forms, implicating them in early synaptic deficits (*1*). Despite this well-described toxicity, Aβ is present in the brains of healthy subjects at low picomolar (pM) concentrations (*2-4*) and is required for normal memory and long-term potentiation (LTP), a type of synaptic plasticity thought to underlie memory formation (*5*). Indeed, brief exposure to pM Aβ enhances memory and LTP (*6, 7*), whereas prolonged exposure impairs them, inducing synaptic alterations such as presynaptic remodeling, a higher basal number of functional presynaptic release sites, and increased spontaneous neurotransmitter release (*6*). Interestingly, aging, the greatest risk factor for sporadic AD, may contribute to this process by reducing neprilysin, an enzyme involved in Aβ clearance in humans and rodents (*8, 9*), thereby prolonging Aβ presence during neural activity (*10, 11*). These findings raise a fundamental question: how does Aβ shift from a physiological modulator to a pathological agent in AD?

To address this, we analyzed the epigenomics and transcriptomics changes induced by short and prolonged exposure to pM Aβ_42_ in wild-type mice, with a particular focus on DNA methylation, a mechanism increasingly linked to AD pathogenesis (*12-14*). Previous studies suggest that Aβ can directly alter DNA methylation (*15, 16*), and our findings support this, revealing a time-dependent Aβ-driven modulation of gene methylation and expression. Notably, prolonged exposure alters methylation and expression of genes involved in synaptic function (*6*), supporting the view that abnormal synaptic function and memory occur as a consequence of prolonged exposure to Aβ at pM doses. Interestingly, short exposure to Aβ at pM doses increased the expression of genes related not only to synaptic function and memory, but also to immune responses, aligning with evidence of Aβ’s antimicrobial properties and its role in innate immunity (*17-20*). Furthermore, we also identified prolonged exposure– dependent regulation of genes involved in calcium (Ca^2+^) homeostasis, and an intermediate ‘transition’ phase marked by intact memory and distinct epigenetic signatures.

Together, these results identify Aβ as an epigenetic regulator and provide mechanistic insights into how prolonged, low-dose exposure may initiate AD pathology. This work offers a new framework for understanding early molecular events in AD and for exploring gene-environment interactions that influence disease onset.

## Results

### Short and long exposures to pM Aβ have opposite effects on contextual memory in mice

To evaluate the impact of Aβ on associative memory, we used the contextual fear conditioning (FC) test, which simulates associative learning memory, a type of memory commonly impaired in AD patients (*21*). Mice received seven intracerebroventricular injections at 20-minute intervals with either pM oligomeric Aβ_42_ (oAβ) or vehicle (Fig. 1A). Experimental groups differed in the timing of Aβ exposure: ‘Vehicle’ received seven vehicle injections; ‘Aβ1’ received six vehicle injections followed by one oAβ injection immediately prior to FC; ‘Aβ4’ received four oAβ injections following three vehicle injections; and ‘Aβ7’ received oAβ in all seven injections. Freezing behavior during cued fear learning control was unaffected across groups, indicating no involvement of amygdala (Fig. S1).

**Fig. 1:**
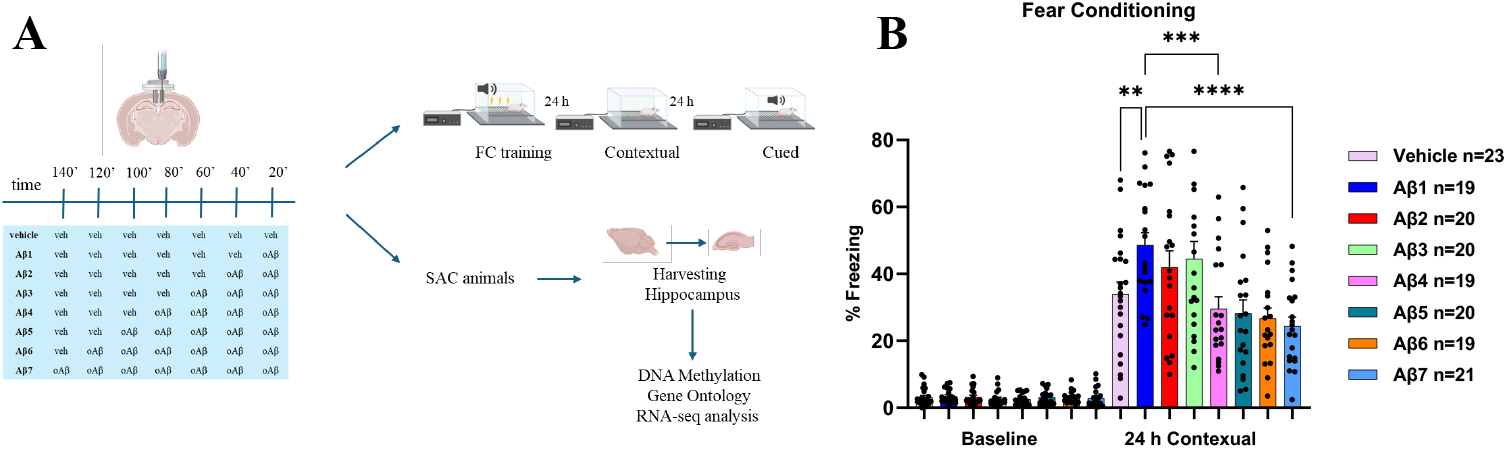
Time-course of the experimental design eliciting associative memory after hippocampal infusion of pM Aβ or vehicle. (A) Schematic showing the experimental design. SAC: sacrifice. Veh: vehicle. Created with BioRender.com released under a Creative Commons Attribution-NonCommercial-NoDerivs license. (B) Contextual fear memory performance is compromised following long exposure of 200 pM Aβ. (Contexual:1-way ANOVA, F (7,153) = 5.232, p < 0.0001, baseline 1-way ANOVA: F (7,153) = 0.3007, p = 0.9526. Nonparametric T-test: vehicle vs Aβ1 p=0.0075 (**), Aβ1 vs Aβ4 p=0.0008 (***), Aβ1 vs Aβ7 p<0.0001 (****).

The results indicated an increase in memory at Aβ1 corresponding to 1 injection of pM oAβ into the hippocampus, 20 min prior to the electric shock. A decline in memory performance started at Aβ4, which corresponds to 4 injections of pM oAβ into the hippocampus, with the first injection of pM oAβ administered 80 minutes before the test (Fig. 1B). This is consistent with the study by Koppensteiner *et al*. showing that mouse hippocampal slices perfused with pM oAβ for 20 minutes significantly increased LTP 1 hour after perfusion, whereas potentiation was reduced after 3-hour perfusion (*6*). These findings suggest that prolonged exposure to pM concentrations of oAβ shifts the peptide’s effect on memory from an increase to detrimental, ultimately impairing associative memory

### pM Aβ has a time-dependent effect on DNA methylation

DNA methylation is one of the most important mechanisms regulating gene expression. To understand if pM doses of oAβ had a time-dependent effect on DNA methylation of murine hippocampi, we first performed a methyl sequencing (methyl-seq) analysis on the samples that were collected immediately after the electric shock. We focused on four main conditions: vehicle, Aβ1, Aβ4 and Aβ7. Aβ1 was selected as the time point associated with an improvement in FC, while Aβ4 and Aβ7 were chosen because they represent, respectively, the point at which associative memory is closest to baseline levels and the stage associated with associative memory decline compared to vehicle. We compared these different conditions to identify which genes were differentially methylated over time.

The heatmaps show the differentially methylated regions (DMRs) identified following short and prolonged exposure to pM Aβ (Fig.s 2A–2C, Table S1). The greatest differences were observed in Aβ1/Aβ4 reflecting changes in the transition phase between early enhanced memory and late memory loss where 261 DMRs were found significantly changed compared, respectively, to the 142 and 204 regions in Aβ4/Aβ7 and vehicle/Aβ1 indicating that exposure of Aβ for a prolonged time has a remarkably different effect on DNA methylation. In addition, 256 DMRs were identified in Aβ1/Aβ7 (Fig. S2). This supports the observation that short and prolonged exposure to Aβ induce different changes in DNA methylation. The values for the differentially methylated cytosines (DMCs) can be found in Table S2. Interestingly, we also found that in Aβ1/Aβ4 most of the DMRs, 35 out of 261 regions, were localized in chromosome 8 while in Aβ4/ Aβ7 most of the DMRs, 18 out 142 regions, were localized in chromosome 1, and in vehicle/ Aβ1, 21 out of 204 regions, were localized in chromosome 2 (Fig.s 2D–2F). This suggests that short and long exposure to pM oAβ are associated with methylation of genes localized in different chromosomes. Furthermore, short exposure to Aβ is associated with a high level of hypomethylation (pink in the circos plot) while prolonged exposure with hypermethylation (green in the circos plot) (Fig. 2D-F).

**Fig. 2:**
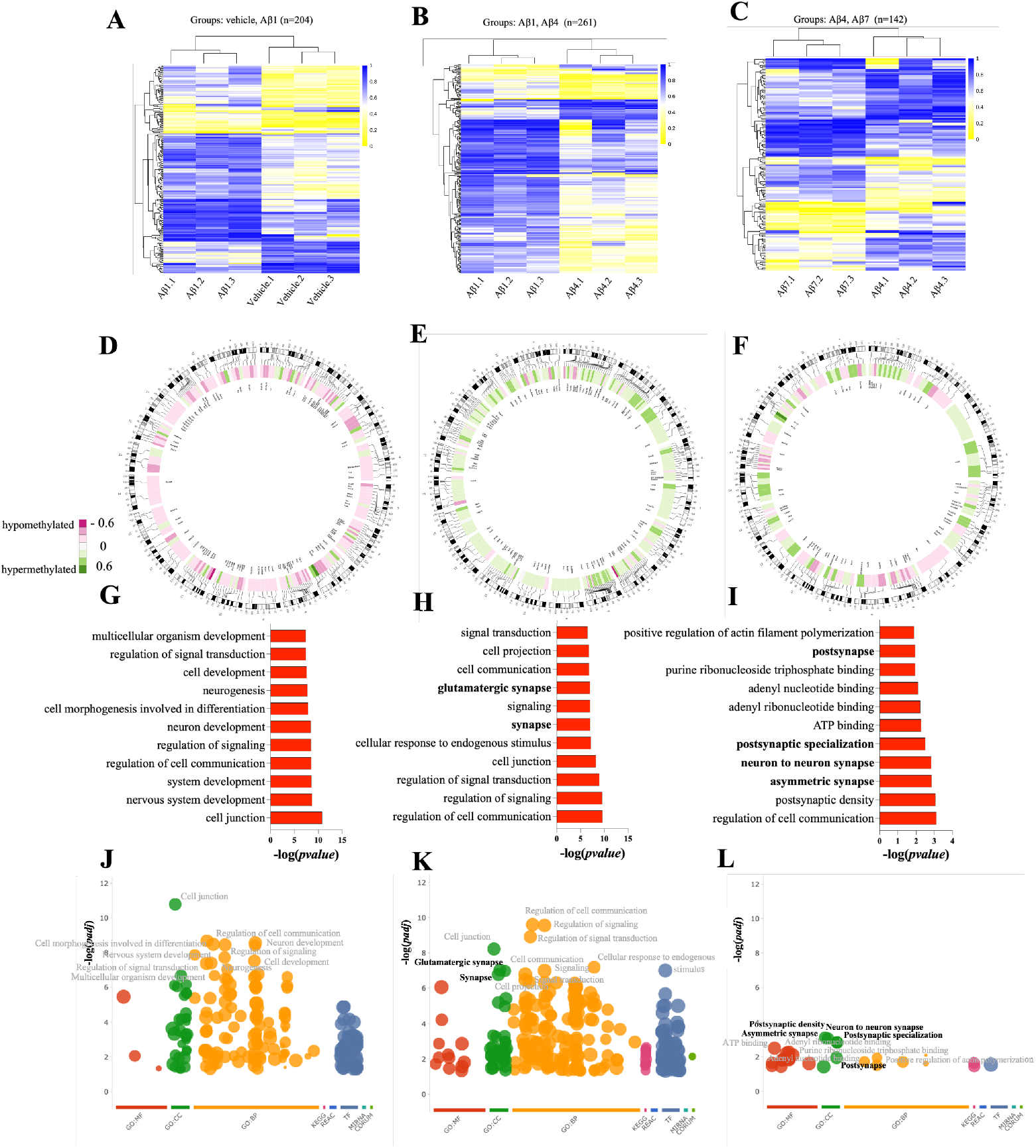
pM oAβ has a time-dependent effect on DNA methylation. (A-C) Cluster heatmaps of DMRs for Vehicle/Aβ1 (A), Aβ1/Aβ4 (B), Aβ4/Aβ7 (C) (FDR ≤ 0.05 and absolute methylation difference ≥ 0.1). (D-F) Circos plots representing chromosomes associated with the differentially methylated regions (DMRs) identified: Vehicle/Aβ1 (D), Aβ1/Aβ4 (E), Aβ4/Aβ7 (F). Circular tracks from outside to inside indicate chromosome number and length, DMR location, cluster heatmap showing the difference in methylation level, and names of genes corresponding to the DMRs. (G-I) GO enrichment analysis of DMRs for Vehicle/Aβ1 (G), Aβ1/Aβ4 (H), Aβ4/Aβ7 (I). Pathways are shown in ascending order based on –log_10_ (Pvalue). (J-L) Manhattan plots for Vehicle/Aβ1 (J), Aβ1/Aβ4 (K), Aβ4/Aβ7 (L). The x-axis represents functional terms of DMRs that are grouped and color-coded by data sources (e.g. Molecular Function from GO is red). The y-axis shows the adjusted enrichment p-values in negative log10 scale. GO terms related to the synapse are highlighted in bold letters.

### Gene Ontology analysis of methylated genes

Genes overlapping with DMRs were subjected to Gene Ontology (GO) analysis to assess functional enrichment. In vehicle/Aβ1, ‘nervous system development’ emerged as one of the most enriched terms. This is consistent with the study by Zhou *et al*. showing that modest concentrations of Aβ play a key role in synaptogenesis (*22*). The genes that were differentially methylated in Aβ1/Aβ4 reflecting changes in the transition phase were enriched on biological pathways involved in ‘synapses’ as shown in the bar graphs and Manhattan plot (Fig.s 2H and 2K). This suggests that pM Aβ regulates the methylation of genes involved in the synapses and that this regulation is time-dependent. Gene ontology (GO) analysis also revealed that genes related to the synapses were enriched in Aβ4/Aβ7 supporting the hypothesis that prolonged exposure to pM oAβ plays a central role in modulating the methylation of synaptic genes (Fig.s 2I and 2L). This is also supported by the GO analysis of vehicle/Aβ7, which identified ‘synapse’ and ‘glutamatergic synapse’ as the two most significantly enriched terms (Fig. S2). Interestingly, genes related to the ‘post synapses’ and ‘postsynaptic density’ were identified in Aβ4/Aβ7 indicating that pM Aβ could play a role in the methylation of distinct genes related to synaptic function at different times. Furthermore, biological pathways involved in ‘signal transduction’ and ‘signaling’ were found enriched both in vehicle/Aβ1 and Aβ1/Aβ4 but not in Aβ4/Aβ7, suggesting that Aβ has a time-dependent effect on the methylation of genes related to signaling (Fig.s 2G–2L).

### Gene expression is affected by pM Aβ at different time points

To understand the effect of pM oAβ on the gene expression, we performed RNA sequencing (RNA-seq) analysis. Table S3 shows the normalized TPM values in the conditions of interest. To correlate the results from the RNA-seq with the data from the Methyl-seq, we performed an integrated analysis as shown in the Venn diagrams (Fig.s 3A–3C and S3). Fig. 4A shows the heatmap with the TPM (transcripts per million) values of the 52 genes that were found both differentially expressed, DEG (differentially expressed genes), and methylated in at least one condition. Table S4 describes the fold change and p-value of the genes identified in the different conditions. Nominal p-values were used in the DEG analysis. To reduce false positives, we performed an ANOVA on the 52 genes of interest (Table S5).

**Fig. 3:**
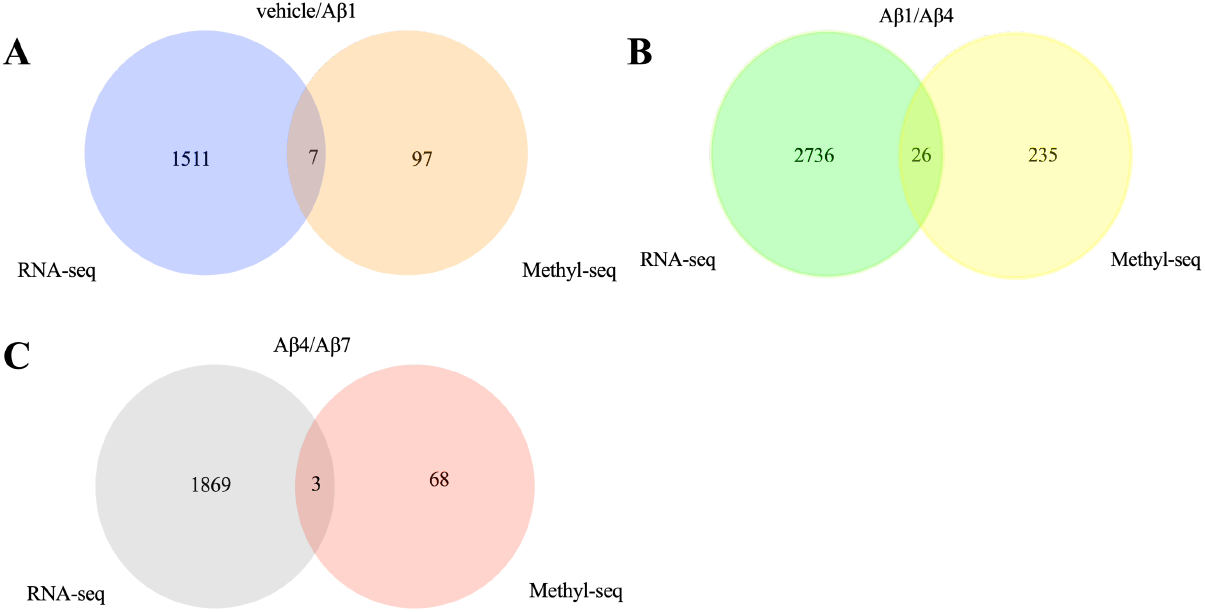
Integrated analysis of DMRs and DEGs following exposure to pM Aβ. (A-C) Venn diagrams representing the overlap of DEGs and DMRs in vehicle/ Aβ1 (A), Aβ1/Aβ4 (B), Aβ4/Aβ7 (C). DEGs p ≤ 0.1.

**Fig. 4:**
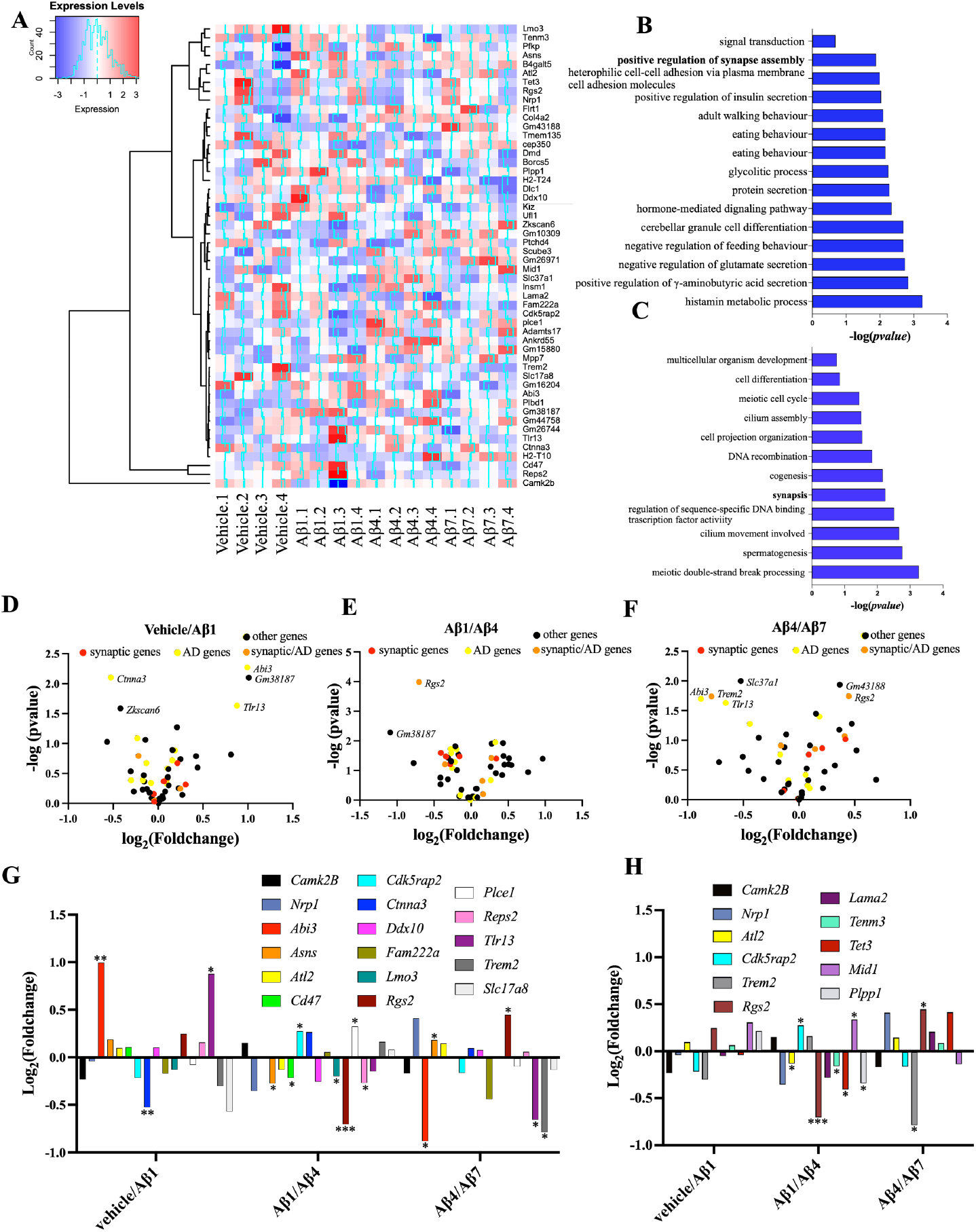
Changes in RNA expression in genes that are differentially methylated following exposure to pM Aβ. (A) Cluster heatmap showing the expression level of the genes found differentially methylated and expressed in at least one condition for vehicle, Aβ1, Aβ4, and Aβ7. p ≤ 0.1. (B-C) GO enrichment analysis for Aβ1/Aβ4 (B) and Vehicle/Aβ1 (C). Pathways are shown in ascending order based on –log(pvalue). (D-F) Scatter plots of differentially expressed genes (DEGs) for vehicle/Aβ1 (D), Aβ1/Aβ4 (E), and Aβ4/Aβ7 (F). Genes are classified into three categories based on their role in synapses (red), AD (yellow), or both (orange). Names of genes with -log(pvalue) ≥1.5 are shown. (G-H) Histogram representing the DEGs associated with AD (G) and synaptic function (H) found differentially methylated and expressed in at least one condition. *, p ≤ 0.05, **, p ≤ 0.01; ***, p ≤ 0.001.

Most of the DMRs identified were localized in the gene bodies and hypermethylated in Aβ1/Aβ4 and Aβ4/Aβ7 whereas they were hypomethylated in vehicle/Aβ1 (Tables S1). While hypermethylation of CpG islands in the promoter regions silences transcription of genes, hypermethylation within gene bodies can be associated with either increase or decrease in gene expression (*23-25*). We observed both upregulation and downregulation of gene expression.

### Genes involved in the synapses and immune response are differentially expressed and methylated after exposure to pM Aβ

GO analysis for the RNA-seq data indicates that DEGs were enriched on biological pathways involved in ‘synapses’ in Aβ1/Aβ4 (Fig. 4B). This result was consistent with the previous GO analysis on the Methyl-seq data and indicates that prolonged exposure to pM Aβ affects both DNA methylation and gene expression of synaptic genes. The only significantly enriched term related to the synapses in vehicle/Aβ1 reflecting changes in the early enhanced memory phase was the ‘positive regulation of synaptic assembly’ (Fig. 4). 11 out of the 52 genes identified to be differentially expressed and methylated in at least one condition of interest were involved in synaptic function. Some of these genes had a distinct expression pattern. They were downregulated in Aβ1/Aβ4 reflecting changes in the transition phase and upregulated in Aβ4/Aβ7 reflecting changes in the late memory impaired phase suggesting that pM Aβ can modulate the expression of the same genes over time (Fig.s 4D–4H). Among the genes associated with synaptic function identified, *Nrp1, Rgs2*, and *Trem2* have been associated with AD (*26-28*). *Rgs2* is pivotal for post- and pre-synaptic function(*29*), and it was found to be downregulated in patients with AD. This is consistent with our experiment showing that *Rgs2* is downregulated in Aβ1/Aβ4 and suggests that as exposure to pM Aβ starts to be protracted with time, its levels are downregulated. Interestingly, *Mid1*, a gene encoding an ubiquitin E3 that catalyzes the degradation of PP2A, a phosphatase known to be reduced in AD (*30*), is upregulated in Aβ1/Aβ4 (*31*). We also identified differentially methylated and expressed genes, such as *Tenm3* and *Tet3*, which have not been previously linked to AD but are involved in synaptic function(*32, 33*). Given the importance of synaptic dysfunction in AD, we created scatter plots for the conditions of interest that highlight the genes involved in synaptic function and those that have already been associated with AD (Fig.s 4D-4F).

Among genes previously implicated in AD, *Abi3* and *Tlr13* exhibited similar expression patterns. They were upregulated in vehicle/Aβ1 and downregulated in Aβ4/Aβ7 (Fig. 4G). *Abi3* and *Tlr13* play both a role in the innate immune response indicating that Aβ might support the immune response (*34, 35*). In addition, *Tlr2*, a member of the TLR family, was upregulated in vehicle/Aβ1 (Fig. S6). This might indicate a correlation between pM Aβ and TLR expression and is supported by studies showing that Aβ stimulates *Tlr2* expression in microglia (*36*). The role of Aβ in the immune response is further supported by the observation that genes found upregulated in vehicle/Aβ1, including *Cd14, Trem1, Cd33*, and *EphA1* are involved in the innate immune response. Interestingly, some of these genes have been linked to AD (*37-39*).

### Prolonged exposure to pM Aβ affects the expression of genes involved in Ca^2+^homeostasis

Koppensteiner *et al*. demonstrated that prolonged exposure to pM Aβ reduced glutamate-induced plasticity in primary hippocampal neurons while short exposure increased it (*6*). This time-dependent impairment induced by pM Aβ was also described on LTP in hippocampal slices (*2*). Ca^2+^ signaling is critical in synaptic plasticity, and its disruption has been associated with AD (*40-42*). In addition, it has been described that Aβ peptides destabilize neuronal Ca^2+^ regulation in cortical neurons (*43*). LTP is triggered by the elevation of intracellular Ca^2+^ levels controlled by glutamate receptors, transient receptors potential (TRP), and voltage-gated calcium channels. Therefore, we analyzed the expression pattern of these receptors and channels, to understand if prolonged exposure to pM oAβ could induce their up or downregulation (Fig. 5A).

**Fig. 5:**
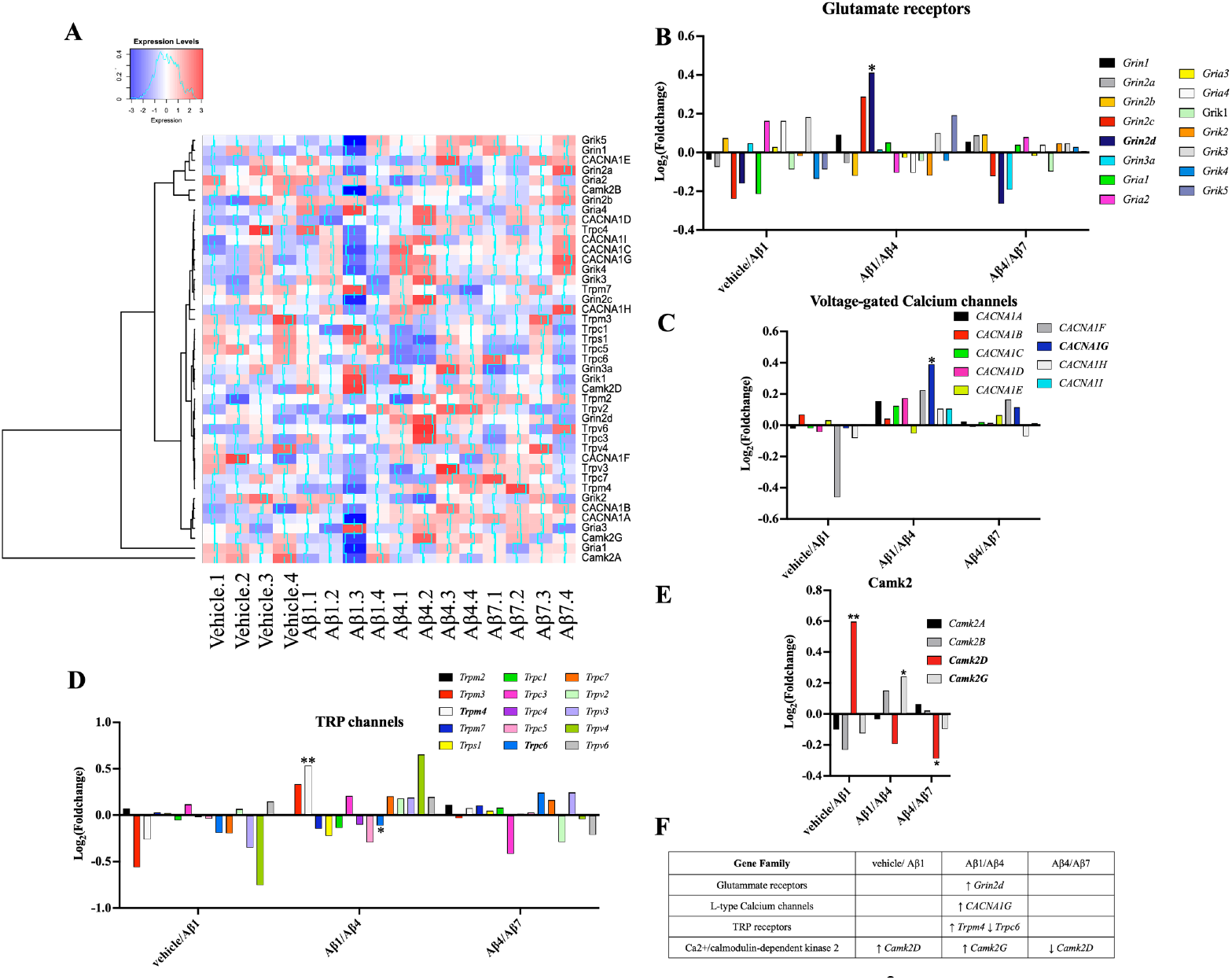
Changes in RNA expression in genes involved in Ca^2+^ homeostasis following exposure to pM Aβ. (A) Cluster heatmap showing the expression level of the genes involved in Ca^2+^ homeostasis. (B-E) Histogram representing the genes encoding glutamate receptors (B), Voltage-gated calcium channels (C), TRP channels (D), and Camk2 (E). *, p ≤ 0.05, **, p ≤ 0.01 (F), Table summarizing the DEGs identified in Vehicle/Aβ1, Aβ1/Aβ4, and Aβ4/Aβ7.

**Fig. 6:**
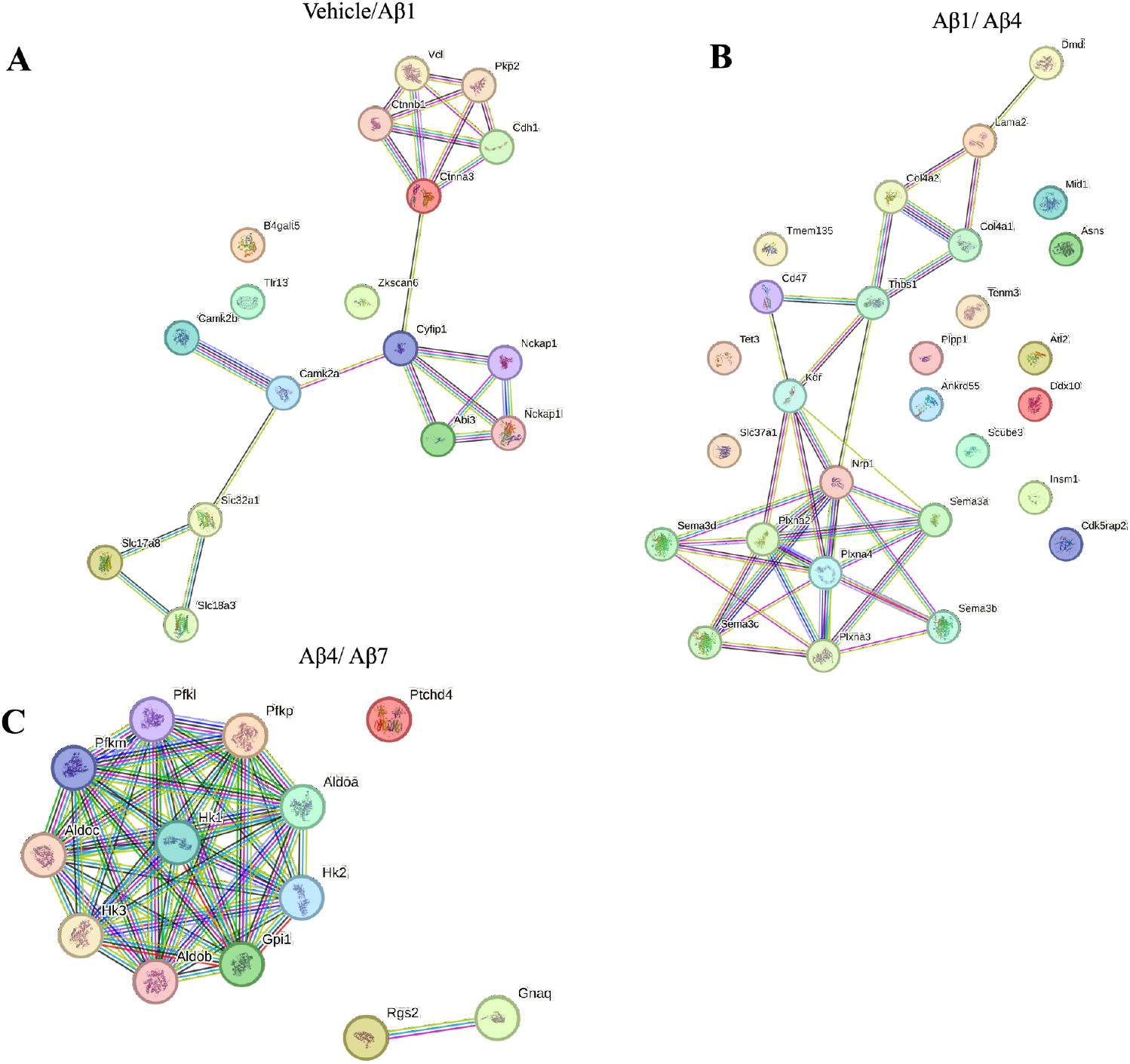
Pathways analysis of differentially methylated genes following exposure to pM Aβ. (A-C) STRING analysis was performed on the genes found differentially methylated and expressed in vehicle/Aβ1(A), Aβ1/Aβ4 (B) and Aβ4/Aβ7 (C). For the analysis, text mining, experiments, and databases were chosen for active interaction sources. A value of 0.400 was selected as the minimum required interaction score. For the analysis, text mining, experiments, and databases were chosen for active interaction sources, and a value of 0.400 was selected as the minimum required interaction score. Line colors represent known interactions from curated databases (blue), experimentation (purple), gene neighborhood (green), gene fusions (red), gene co-occurrence (blue), text mining (yellow), and co-expression (black).

Fig. 5A shows the heatmap with the TPM values of the genes encoding the proteins of interest. The three main classes of ionotropic glutamate receptors are the NMDA (*Grin*), AMPA (*Gria*), and Kainate (*Grik*) (Fig. 5B). *Grin2d* was the only gene of this family upregulated. *Grin2d* encodes the GluN2D subunit of the NMDA receptor that has been found to be associated with LTP in the mouse hippocampus(*44, 45*). Interestingly, the GO analysis for the RNA-seq revealed ‘negative regulation of glutamate secretion’ as a significantly enriched term in vehicle/Aβ4 suggesting that prolonged exposure to pM Aβ might also affect the secretion of glutamate (Fig. S4A). In addition, among the voltage-gated calcium channels, *CACNA1G* was upregulated in Aβ1/Aβ4 (Fig. 5C). *CACNA1G* encodes the Cav3.1 T-type channel. The blockade of Cav3.1 has been shown to impair LTP at parallel fiber-Purkinje cell synapses (*46*). Interestingly, *Trpm4*, a channel that belongs to the TRP family, was also upregulated in Aβ1/Aβ4, vehicle/Aβ7 and Aβ1/Aβ7 (Fig. 5D and S5C). *Trpm4* is a non-selective monovalent cation channel activated by intracellular Ca^2+^ during excitatory responses in hippocampal neurons and a co-activator of NMDA receptors during the induction of LTP (*47*), (*48, 49*). Therefore, upregulation of *Trpm4* may impair LTP by depolarizing the membrane, inactivating voltage-gated Ca^2+^ channels, and reducing intracellular Ca^2+^ influx.

The Methyl-seq analysis revealed that *Camk2b* is differentially methylated across all the conditions of interest. Interestingly, we identified two distinct DMRs in vehicle/Aβ1 while only a single DMR was present in the other conditions (Table S1). The product of the *Camk2b* gene belongs to the Ca^2+^/calmodulin-dependent protein kinase 2 (Camk2) family and is one of the major regulators of neurotransmitter release associated with AD (*50*). The Camk2 family consists of four different isozymes CAMK2A, CAMK2B, CAMK2C, and CAMK2D(*51*).

Interestingly, we did not find *Camk2b* significantly differentially expressed. However, we found that *Camk2d* was upregulated in vehicle/Aβ1 and downregulated in Aβ1/Aβ7 (Fig. 5E). CAMK2D is less studied than its paralogs CAMK2KB and CAMK2A, which are known to be required to produce LTP and are the predominant isoforms in the hippocampus (*52*). Our result suggests that this isoform might play a role in the time-dependent effect on LTP mediated by pM Aβ.

Our findings suggest that pM Aβ is associated with the differential expression of distinct genes involved in memory formation and Ca^2+^ homeostasis at different time points. The most significant changes, including the upregulation of *Trpm4*, were observed in Aβ1/Aβ4 (Fig. 5F). This is aligned with our FC results (Fig. 1B) and the previous LTP studies (*6*). Interestingly, these genes encode proteins that are known to be localized postsynaptically. However, our integrated analysis also identified *Cdk5rap2* (Fig. 4G), a gene differentially expressed and methylated in Aβ1/Aβ4, which encodes for CDK5RAP2, an interactor of the CDK5 activator, CDK5R1 (*53*). CDK5 is known to contribute to Ca^2+^ homeostasis and neurotransmitter release (*54*). In addition, *Rgs2* has been identified to be important in controlling presynaptic Ca^2+^ channels (*29*). This suggests that pM Aβ might disrupt both pre- and post-synaptic signaling.

### Protein-protein interaction (PPI) interaction networks are induced by pM Aβ

In order to understand the interactions among the proteins encoded by the genes found differentially methylated and expressed, a protein-protein interaction (PPI) analysis was performed using Search Tool for the Retrieval of INteracting Genes/ Proteins (STRING). Of the proteins encoded by the genes that were found both differentially methylated and expressed in vehicle/Aβ1 (Fig. 4A and Table S4), 4 proteins clustered in a large network and 3 proteins were not clustering (Fig. 5A). Interestingly, the network included NKCAP1I. *Nckap1I* was not found to be differentially methylated but was upregulated in the RNA-seq analysis. This could suggest that the methylation of genes in this pathway could have a direct effect on the expression of *Nckap1l*. NKCAP1L is a regulator of the actin cytoskeleton and, together with ABI3, is a component of the WAVE complex (*55*). In Aβ1/Aβ4, 6 proteins were clustering and 13 were not clustering (Fig. 5B). Similarly to *Nckap1l, Plxna3* was not found to be differentially methylated but was differentially expressed. PLXNA3 is a protein that plays a role in neuronal development (*56*). Among the proteins clustering in this network, CD47 and NRP1 are both involved in AD (*28, 57*). In Aβ4/Aβ7, PFKB clustered in a large network while RGS2 clustered in a small network (Fig. 5C). The STRING analysis shows that PFKB interacts with HK1, a hexokinase that was reported to be differentially expressed in neuronal progenitor cells of patients with AD (*58*). This suggests that prolonged exposure to pM Aβ could disrupt the cell energy metabolism.

Interestingly, 46 out of 103 genes upregulated in vehicle/Aβ1 with a log_2_(fold change) ≥1 and p≤0.05 were associated with the immune response, with the majority involved in innate immunity and inflammation. By performing a PPI analysis through the STRING database, we observed that the proteins encoded by these genes clustered in a large network (Fig. S6). Similarly, proteins encoded by genes involved in the immune response and upregulated in vehicle/Aβ4 and vehicle/Aβ7 formed a network (Fig. S7 and S8). IL6 and IL11, interleukins known for their pro-inflammatory roles, were at the center of the network for both conditions. IL6 has been linked to cognitive impairment in AD (*59*). This is consistent with the GO analysis for vehicle/Aβ7 showing the ‘cellular response to IL6’ as one of the most significantly enriched terms. Interestingly, only some of these genes, such as *Lilrb4a*, a gene encoding for a protein mediating suppressive immune responses, overlapped with those identified in vehicle/Aβ1 (*60*). The majority were distinct.

We did not observe an interaction network for upregulated genes in Aβ1/Aβ7. However, PPI analysis of proteins encoded by downregulated genes revealed a network with MMP-9, a protein critical for memory formation, at its center (Fig. S9) (*61*). MMP-9 was also downregulated in vehicle/Aβ7 and Aβ4/Aβ7 (Table S4), suggesting that prolonged exposure to pM Aβ strongly affects its expression. MMP-9 is required for hippocampal LTP in mice (*61*). This is consistent with our previous studies showing that prolonged exposure to pM Aβ impairs LTP (*6*).

## Discussion

How Aβ transitions from a physiological to a pathological agent in the brain remains poorly understood. Here, we demonstrate that the decline in associative memory and synaptic plasticity previously observed following prolonged exposure to pM Aβ in wild-type mice (*6*) is mediated by alterations in DNA methylation and gene expression. Our integrative analysis identified time-dependent changes in genes regulating synaptic function, immune signaling, and calcium homeostasis, providing new insights into the molecular events initiating AD.

Prolonged Aβ exposure disrupted methylation and expression of genes linked to synaptic function and AD, including *Trem2* and *Rgs2* (*27, 62*). *Rgs2*, which modulates presynaptic Ca^2+^ channels and short-term synaptic plasticity (*29, 63*), was downregulated at later stages of Aβ exposure. Given the critical role of calcium homeostasis in memory and synaptic signaling (*40*), we examined other components of this pathway. *Camk2b*, encoding a major calcium-responsive kinase involved in synaptic plasticity (*64*), was differentially methylated across all Aβ exposure stages. Furthermore, genes controlling Ca^2+^ influx—*Grin2d, CACNA1G*, and *Trpm4*—were upregulated with prolonged Aβ exposure. Importantly, downregulation of *Trem2*, which exacerbates neuroinflammation in AD models (*65*), also emerged as a signature of chronic Aβ exposure. Together, these findings suggest that prolonged exposure to pM Aβ dysregulates Ca^2+^-dependent synaptic gene programs through dynamic epigenetic changes.

This dynamic role of Aβ was also evident in the regulation of genes involved in the immune response (*34, 35*). *Abi3* and *Tlr13* were upregulated after short Aβ exposure but downregulated after prolonged exposure, supporting previous studies showing that Aβ exerts antimicrobial activity by modulating the immune response (*17*). Our data also reveal that short exposure to Aβ increases the expression of *Tlr2*, suggesting that Aβ may act as a master regulator of TLR receptors expression. While Tlr13 was both differentially expressed and methylated, *Tlr2* was found to be only upregulated, indicating that methylation might trigger a cascade effect leading to overexpression of genes within the same pathways. Interestingly, IL6 and IL11, which have been linked to AD and cognitive decline (*59, 66*), were at the center of the network in the PPI analysis for vehicle/Aβ7 and vehicle/Aβ4. These findings align with studies showing IL6 impairs neurotransmitter release and LTP, implying that Aβ-driven immune gene expression may contribute to synaptic dysfunction over time (*67, 68*).

The data suggest that Aβ functions as an epigenomic regulator. The role of Aβ in regulating DNA methylation has been previously described (*15, 16*), our study highlights the time-dependent nature of these changes in relation to the memory and synaptic impairment (*6, 7*), providing insights into the specific genes and pathways affected. Many of the upregulated genes identified when comparing the vehicle to the Aβ treatment encoded for proteins involved in the immune response. The STRING revealed that these proteins formed large networks. We also identified a large network of proteins encoded by genes downregulated in Aβ1/Aβ7. Interestingly, MMP-9 was at the center of this network, suggesting that prolonged exposure to pM Aβ significantly regulates its expression. MMP-9 plays a role in synaptic plasticity, as hippocampal LTP and FC have been described to be impaired in MMP-9 null-mutant mice (*61*). In addition, it has been found to be crucial for Aβ40 and Aβ42 degradation (*69*). This suggests that *Mmp9* downregulation might contribute to the associated memory decline observed after prolonged exposure to Aβ pM.

These findings provide insight into the causal relationship between prolonged exposure to pM oAβ and the impairment of memory formation and synaptic plasticity, finding its origins in Aβ-induced alterations in DNA methylation and RNA expression. Furthermore, these results shed light on the mechanism by which Aβ shifts from having a memory enhancing effect to a toxic one. Many genes associated with LTP impairment, such as *Trem2, Rgs2, CACNA1G, Il6, Il11*, and *Mmp9*, were differentially expressed following prolonged exposure to pM Aβ but not following short exposure. Interestingly, we identified *Cam2kd* upregulated after short exposure. *Camk2d* encodes for CAMK2D, a paralog of CAMK2A. The overexpression of CAMK2A has been found to potentiate synaptic transmission (*70*). Although CAMK2D is less abundant than its paralogs CAMK2A and CAMK2B, it plays a crucial role in long-term memory in the mouse hippocampus. Notably, its mRNA expression remains elevated up to a week after training (*71*). Our results suggest that the effect described by Zalcam *et al*. may underlie the beneficial impact of short Aβ exposure.

One of the limitations of this study is that our experimental design cannot be translated into examining the progressive development of AD in patients. Brain tissues can only be collected postmortem. Patients diagnosed with AD usually have molecular and cellular changes decades before symptoms, such as severe memory deficits. Therefore, it is not possible to perform this type of study in humans. Nevertheless, our findings offer valuable insights into early epigenomic alterations linked to memory decline that may also occur in human disease.

In our analysis, we prioritized biological signals over conservative statistical thresholds to enhance sensitivity. DEGs were identified using nominal p-values (p < 0.05), without correction for multiple testing. This approach was intentionally chosen to maximize the inclusion of potentially relevant genes for downstream integrative analysis with DNA methylation data. To further evaluate expression differences among the identified DEGs, we performed an ANOVA to assess variation across experimental conditions. We acknowledge that the lack of multiple testing correction may increase the rate of false positives, which should be considered when interpreting the findings.

This study lays the foundation to unravel the origins of AD. We identified key genes involved in the transition of Aβ from being beneficial to having a toxic effect. Our data reveal that Aβ induces a complex epigenetic effect that leads to the decline of associative memory. Many of the DEGs identified were linked to synaptic function and Ca^2+^ homeostasis (see summary in Fig. S10). The genetic factors identified in this study provide insights into the underlying mechanism of AD pathogenesis and could pave the way for the development of novel therapeutic strategies. Therapeutic approaches, including gene editing tools like CRISPR/Cas9 (*72, 73*), could be employed to correct the aberrant changes observed following prolonged exposure to pM Aβ.

## Acknowledgments

We would like to thank you Dr. Luana Fioriti (Mario Negri Institute for Pharmacological Research IRCCS, Milano, Italy), Dr. Luciano D’Adamio (Rutgers University, Newark, New Jersey, USA), and Dr. Peter Koppensteiner for their comments on the manuscript.

## Author contributions

Methodology: AP, EA, OA, AFT Formal analysis: AP, EA Conceptualization: AP, EA, AS Funding acquisition: OA

Writing – original draft: AP, EA, OA Project administration: OA

Writing – review & editing: AP, EA, OA, AFT

## Data and materials availability

All data are available in the main text or the supplementary materials.

## Supplemental Materials

Materials and Methods Fig. S1 to S10.

Table S1 to S6.

## References

1. D. J. Selkoe, Alzheimer’s disease is a synaptic failure. Science 298, 789–791 (2002).

2. D. Puzzo et al., Picomolar amyloid-beta positively modulates synaptic plasticity and memory in hippocampus. J Neurosci 28, 14537–14545 (2008).

3. P. Mastrangelo et al., Dissociated phenotypes in presenilin transgenic mice define functionally distinct gamma-secretases. Proc Natl Acad Sci U S A 102, 8972–8977 (2005).

4. V. Giedraitis et al., The normal equilibrium between CSF and plasma amyloid beta levels is disrupted in Alzheimer’s disease. Neurosci Lett 427, 127–131 (2007).

5. T. V. Bliss, G. L. Collingridge, A synaptic model of memory: long-term potentiation in the hippocampus. Nature 361, 31–39 (1993).

6. P. Koppensteiner et al., Time-dependent reversal of synaptic plasticity induced by physiological concentrations of oligomeric Abeta42: an early index of Alzheimer’s disease. Sci Rep 6, 32553 (2016).

7. D. Puzzo et al., Endogenous amyloid-beta is necessary for hippocampal synaptic plasticity and memory. Ann Neurol 69, 819–830 (2011).

8. A. Caccamo, S. Oddo, M. C. Sugarman, Y. Akbari, F. M. LaFerla, Age- and region-dependent alterations in Abeta-degrading enzymes: implications for Abeta-induced disorders. Neurobiol Aging 26, 645–654 (2005).

9. N. Iwata, Y. Takaki, S. Fukami, S. Tsubuki, T. C. Saido, Region-specific reduction of A beta-degrading endopeptidase, neprilysin, in mouse hippocampus upon aging. J Neurosci Res 70, 493–500 (2002).

10. J. R. Cirrito et al., Synaptic activity regulates interstitial fluid amyloid-beta levels in vivo. Neuron 48, 913–922 (2005).

11. F. Kamenetz et al., APP processing and synaptic function. Neuron 37, 925–937 (2003).

12. H. Lu, X. Liu, Y. Deng, H. Qing, DNA methylation, a hand behind neurodegenerative diseases. Front Aging Neurosci 5, 85 (2013).

13. K. Lunnon, J. Mill, Epigenetic studies in Alzheimer’s disease: current findings, caveats, and considerations for future studies. Am J Med Genet B Neuropsychiatr Genet 162B, 789–799 (2013).

14. A. S. Yokoyama, J. C. Rutledge, V. Medici, DNA methylation alterations in Alzheimer’s disease. Environ Epigenet 3, dvx008 (2017).

15. K. L. Chen et al., The epigenetic effects of amyloid-beta(1-40) on global DNA and neprilysin genes in murine cerebral endothelial cells. Biochem Biophys Res Commun 378, 57–61 (2009).

16. N. Taher et al., Amyloid-beta alters the DNA methylation status of cell-fate genes in an Alzheimer’s disease model. J Alzheimers Dis 38, 831–844 (2014).

17. D. K. Kumar et al., Amyloid-beta peptide protects against microbial infection in mouse and worm models of Alzheimer’s disease. Sci Transl Med 8, 340ra372 (2016).

18. M. Jorfi, A. Maaser-Hecker, R. E. Tanzi, The neuroimmune axis of Alzheimer’s disease. Genome Med 15, 6 (2023).

19. R. D. Moir, R. Lathe, R. E. Tanzi, The antimicrobial protection hypothesis of Alzheimer’s disease. Alzheimers Dement 14, 1602–1614 (2018).

20. D. K. Kumar, W. A. Eimer, R. E. Tanzi, R. D. Moir, Alzheimer’s disease: the potential therapeutic role of the natural antibiotic amyloid-beta peptide. Neurodegener Dis Manag 6, 345–348 (2016).

21. R. Swainson et al., Early detection and differential diagnosis of Alzheimer’s disease and depression with neuropsychological tasks. Dement Geriatr Cogn Disord 12, 265–280 (2001).

22. B. Zhou, J. G. Lu, A. Siddu, M. Wernig, T. C. Sudhof, Synaptogenic effect of APP-Swedish mutation in familial Alzheimer’s disease. Sci Transl Med 14, eabn9380 (2022).

23. M. M. Suzuki, A. Bird, DNA methylation landscapes: provocative insights from epigenomics. Nat Rev Genet 9, 465–476 (2008).

24. P. A. Jones, Functions of DNA methylation: islands, start sites, gene bodies and beyond. Nat Rev Genet 13, 484–492 (2012).

25. M. P. Ball et al., Targeted and genome-scale strategies reveal gene-body methylation signatures in human cells. Nat Biotechnol 27, 361–368 (2009).

26. A. Hadar et al., RGS2 expression predicts amyloid-beta sensitivity, MCI and Alzheimer’s disease: genome-wide transcriptomic profiling and bioinformatics data mining. Transl Psychiatry 6, e909 (2016).

27. Y. Zhou et al., Human and mouse single-nucleus transcriptomics reveal TREM2-dependent and TREM2-independent cellular responses in Alzheimer’s disease. Nat Med 26, 131–142 (2020).

28. K. H. Lim, S. Yang, S. H. Kim, J. Y. Joo, Identifying New COVID-19 Receptor Neuropilin-1 in Severe Alzheimer’s Disease Patients Group Brain Using Genome-Wide Association Study Approach. Front Genet 12, (2021).

29. J. Han et al., RGS2 determines short-term synaptic plasticity in hippocampal neurons by regulating Gi/o-mediated inhibition of presynaptic Ca2+ channels. Neuron 51, 575–586 (2006).

30. J. M. Sontag, E. Sontag, Protein phosphatase 2A dysfunction in Alzheimer’s disease. Front Mol Neurosci 7, 16 (2014).

31. H. Du et al., MID1 catalyzes the ubiquitination of protein phosphatase 2A and mutations within its Bbox1 domain disrupt polyubiquitination of alpha4 but not of PP2Ac. PLoS One 9, e107428 (2014).

32. H. Yu et al., Tet3 regulates synaptic transmission and homeostatic plasticity via DNA oxidation and repair. Nat Neurosci 18, 836–843 (2015).

33. X. Zhang, X. Chen, D. Matus, T. C. Sudhof, Reconstitution of synaptic junctions orchestrated by teneurin-latrophilin complexes. Science 387, 322–329 (2025).

34. R. Medzhitov, Toll-like receptors and innate immunity. Nat Rev Immunol 1, 135–145 (2001).

35. R. Sims et al., Rare coding variants in PLCG2, ABI3, and TREM2 implicate microglial-mediated innate immunity in Alzheimer’s disease. Nat Genet 49, 1373–1384 (2017).

36. K. L. Richard, M. Filali, P. Prefontaine, S. Rivest, Toll-like receptor 2 acts as a natural innate immune receptor to clear amyloid beta 1-42 and delay the cognitive decline in a mouse model of Alzheimer’s disease. J Neurosci 28, 5784–5793 (2008).

37. K. Fassbender et al., The LPS receptor (CD14) links innate immunity with Alzheimer’s disease. FASEB J 18, 203–205 (2004).

38. P. Hollingworth et al., Common variants at ABCA7, MS4A6A/MS4A4E, EPHA1, CD33 and CD2AP are associated with Alzheimer’s disease. Nat Genet 43, 429–435 (2011).

39. E. N. Wilson et al., TREM1 disrupts myeloid bioenergetics and cognitive function in aging and Alzheimer disease mouse models. Nat Neurosci 27, 873–885 (2024).

40. C. R. Rose, A. Konnerth, Stores not just for storage. intracellular calcium release and synaptic plasticity. Neuron 31, 519–522 (2001).

41. B. C. Tong, A. J. Wu, M. Li, K. H. Cheung, Calcium signaling in Alzheimer’s disease & therapies. Biochim Biophys Acta Mol Cell Res 1865, 1745–1760 (2018).

42. E. K. Webber, M. Fivaz, G. E. Stutzmann, G. Griffioen, Cytosolic calcium: Judge, jury and executioner of neurodegeneration in Alzheimer’s disease and beyond. Alzheimers Dement 19, 3701–3717 (2023).

43. M. P. Mattson et al., beta-Amyloid peptides destabilize calcium homeostasis and render human cortical neurons vulnerable to excitotoxicity. J Neurosci 12, 376–389 (1992).

44. A. V. Eapen et al., Multiple roles of GluN2D-containing NMDA receptors in short-term potentiation and long-term potentiation in mouse hippocampal slices. Neuropharmacology 201, 108833 (2021).

45. D. Liao, N. A. Hessler, R. Malinow, Activation of postsynaptically silent synapses during pairing-induced LTP in CA1 region of hippocampal slice. Nature 375, 400–404 (1995).

46. R. Ly et al., T-type channel blockade impairs long-term potentiation at the parallel fiber-Purkinje cell synapse and cerebellar learning. Proc Natl Acad Sci U S A 110, 20302–20307 (2013).

47. P. Launay et al., TRPM4 is a Ca2+-activated nonselective cation channel mediating cell membrane depolarization. Cell 109, 397–407 (2002).

48. B. C. Fearey et al., A glibenclamide-sensitive TRPM4-mediated component of CA1 excitatory postsynaptic potentials appears in experimental autoimmune encephalomyelitis. Sci Rep 12, 6000 (2022).

49. A. Menigoz et al., TRPM4-dependent post-synaptic depolarization is essential for the induction of NMDA receptor-dependent LTP in CA1 hippocampal neurons. Pflugers Arch 468, 593–607 (2016).

50. Y. J. Wang et al., The expression of calcium/calmodulin-dependent protein kinase II-alpha in the hippocampus of patients with Alzheimer’s disease and its links with AD-related pathology. Brain Res 1031, 101–108 (2005).

51. I. Ninan, O. Arancio, Presynaptic CaMKII is necessary for synaptic plasticity in cultured hippocampal neurons. Neuron 42, 129–141 (2004).

52. A. J. Silva, C. F. Stevens, S. Tonegawa, Y. Wang, Deficient hippocampal long-term potentiation in alpha-calcium-calmodulin kinase II mutant mice. Science 257, 201–206 (1992).

53. X. Wang et al., Identification of a common protein association region in the neuronal Cdk5 activator. J Biol Chem 275, 31763–31769 (2000).

54. B. A. Samuels et al., Cdk5 promotes synaptogenesis by regulating the subcellular distribution of the MAGUK family member CASK. Neuron 56, 823–837 (2007).

55. C. N. Castro et al., NCKAP1L defects lead to a novel syndrome combining immunodeficiency, lymphoproliferation, and hyperinflammation. J Exp Med 217, (2020).

56. R. Oleari et al., PLXNA1 and PLXNA3 cooperate to pattern the nasal axons that guide gonadotropin-releasing hormone neurons. Development 146, (2019).

57. T. Phongpreecha et al., Single-synapse analyses of Alzheimer’s disease implicate pathologic tau, DJ1, CD47, and ApoE. Sci Adv 7, eabk0473 (2021).

58. W. I. Ryu et al., Brain cells derived from Alzheimer’s disease patients have multiple specific innate abnormalities in energy metabolism. Mol Psychiatry 26, 5702–5714 (2021).

59. E. S. N. M. Lyra et al., Pro-inflammatory interleukin-6 signaling links cognitive impairments and peripheral metabolic alterations in Alzheimer’s disease. Transl Psychiatry 11, 251 (2021).

60. H. M. Chen et al., Blocking immunoinhibitory receptor LILRB2 reprograms tumor-associated myeloid cells and promotes antitumor immunity. J Clin Invest 128, 5647–5662 (2018).

61. V. Nagy et al., Matrix metalloproteinase-9 is required for hippocampal late-phase long-term potentiation and memory. J Neurosci 26, 1923–1934 (2006).

62. V. S. Jadhav et al., Trem2 Y38C mutation and loss of Trem2 impairs neuronal synapses in adult mice. Mol Neurodegener 15, 62 (2020).

63. K. J. Gerber, K. E. Squires, J. R. Hepler, Roles for Regulator of G Protein Signaling Proteins in Synaptic Signaling and Plasticity. Mol Pharmacol 89, 273–286 (2016).

64. G. M. van Woerden et al., betaCaMKII controls the direction of plasticity at parallel fiber-Purkinje cell synapses. Nat Neurosci 12, 823–825 (2009).

65. J. B. Ruganzu et al., Downregulation of TREM2 expression exacerbates neuroinflammatory responses through TLR4-mediated MAPK signaling pathway in a transgenic mouse model of Alzheimer’s disease. Mol Immunol 142, 22–36 (2022).

66. D. Galimberti et al., Intrathecal levels of IL-6, IL-11 and LIF in Alzheimer’s disease and frontotemporal lobar degeneration. J Neurol 255, 539–544 (2008).

67. G. D’Arcangelo et al., Interleukin-6 inhibits neurotransmitter release and the spread of excitation in the rat cerebral cortex. Eur J Neurosci 12, 1241–1252 (2000).

68. A. J. Li, T. Katafuchi, S. Oda, T. Hori, Y. Oomura, Interleukin-6 inhibits long-term potentiation in rat hippocampal slices. Brain Res 748, 30–38 (1997).

69. P. Yan et al., Matrix metalloproteinase-9 degrades amyloid-beta fibrils in vitro and compact plaques in situ. J Biol Chem 281, 24566–24574 (2006).

70. D. L. Pettit, S. Perlman, R. Malinow, Potentiated transmission and prevention of further LTP by increased CaMKII activity in postsynaptic hippocampal slice neurons. Science 266, 1881–1885 (1994).

71. G. Zalcman et al., Sustained CaMKII Delta Gene Expression Is Specifically Required for Long-Lasting Memories in Mice. Mol Neurobiol 56, 1437–1450 (2019).

72. L. Cong et al., Multiplex genome engineering using CRISPR/Cas systems. Science 339, 819–823 (2013).

73. P. Mali et al., RNA-guided human genome engineering via Cas9. Science 339, 823–826 (2013).

74. D. Puzzo et al., Amyloid-beta peptide inhibits activation of the nitric oxide/cGMP/cAMP-responsive element-binding protein pathway during hippocampal synaptic plasticity. J Neurosci 25, 6887–6897 (2005).

