## Supplementary material for "Epigenetic Shifts Reveal Alzheimer’s Origins after Sustained Picomolar Aβ Exposure": Materials and Methods; Fig. S1 to S10

### Animals

All the experiments were performed using 3–4 month-old male and female C57BL/6 mice hosted at the Columbia University animal facility. The mice were maintained on a 12 h light/dark cycle in stable conditions in terms of temperature, humidity, and ventilation. Water and food were offered ad libitum.

### Aβ oligomers

Oligomeric $A\beta_{1\text{-}42}$ was prepared as described previously(*74*). Briefly, the lyophilized peptide (American Peptide) was resuspended in 100% 1,1,1,3,3,3-hexafluoro-2-propanol (HFIP; Sigma, St. Louis, MO) to 1 mm. The solution was aliquoted, and the HFIP was allowed to evaporate in the fume hood. The resulting clear peptide film was dried under vacuum in a SpeedVac and stored at -20°C. Twenty-four hours before use, the aliquots were added to dimethylsulfoxide (DMSO; Sigma) and sonicated for 10 min. Oligomeric $A\beta_{1\text{-}42}$ was obtained by diluting $A\beta_{1\text{-}42}$-DMSO into ACSF concentration, vortexed for 30 s, and incubated at 4°C for 24 h. Before use, this compound was added to ACSF to obtain 200 nm. $A\beta_{1\text{-}42}$ with scrambled sequence was diluted in ACSF at 200 nmol and then at 200pM immediately before use. Amyloid β-Protein (1-42) is sold by Bachem (Product Number: 4014447)

### Stereotaxic Surgery and Infusion of Aβ

Before fixing the mice in the stereotaxic device, anesthesia was induced by intraperitoneal (i.p.) injection with Avertin (500 mg/Kg). Animals were injected with analgesic (Carprofen subcutaneously on the back, 5 mg/Kg) and local anesthetic (Marcaine subcutaneously under the scalp 3 mg/Kg). A midline incision was made in the skull and the underlying area was cleared of tissue by using $H_2O_2$. The coordinates of the dorsal hippocampus were 2.46 mm posteriorly and 1.5 mm laterally from Bregma to a depth of 1.30 mm [Paxinos G, Franklin K. Paxinos and Franklin's the mouse brain in stereotaxic coordinates. 4th ed: San Diego, Elsevier Academic Press; 2012]. A 26-gauge guide cannula (PlasticsOne; Roanoke, VA) was fixed to the skull using acrylic dental cement (Paladur). After 6–9 days of recuperation period, awake mice were restrained and injected onto both dorsal hippocampi with oAβ (500 nM, 1 µl per side, at 180 and 20 min prior to the task) or vehicle. For the injections, we utilized Hamilton syringes connected with a polyethylene tube at the end of which an internal microsyringe was fixed. The microsyringe was custom-made to reach a depth of 1.5 mm, ensuring that it was not extending beyond the cannula. Following injection, the microsyringe was left in place for 1 min on each side to ensure perfusion of the hippocampus with oAβ. Correct positioning of the cannulas was verified at the end of the experiment by intrahippocampal injection of methylene blue. Administration of pM Aβ occurred on the first experimental day. Mice were injected with pM Aβ or vehicle 7 times every 20 min prior to the test according with the condition.

### Behavioral Studies

The fear conditioning test was used for evaluating associative fear memory in rodents. The test consists in total of 2 days, in which the first day the animals are placed in the fear conditioning chamber (Noldus) for 2 min before the presentation of the conditional stimulus (tone; 2880 Hz at 85 Db). In the last 2 s of the tone, mice received the unconditional stimulus (foot shock; 0.8 mA). After the pairing of the 2 stimuli, mice were left in the chamber for another 30 s in the absence of a stimulus. The second day, mice were returned to the same conditioning chamber for another 5 min without the presence of tone of shock. Freezing behavior, distinguished by the absence of movement except breathing, was monitored during the test using a vision tracking and analysis system (Ethovision XT, Noldus). Cued fear responses were assessed 48 hours after the shock by exposing the mice to the tone in a novel environment.

**Animal euthanasia and sample collection**

Mice were sacrificed through cervical dislocation immediately after shock exposure in a fear conditioning chamber. Hippocampi were separated from the rest of the brain and frozen in liquid nitrogen and stored in a -80C freezer. The samples were collected for the following analysis.

**Statistical analysis**

For the behavioral tests, animals were run in cohorts in which sex of mice was kept balanced across groups. Results were analyzed with one-way ANOVA with Bonferroni post-hoc correction in FC behavior experiment.

**DNA Methylated analysis: Whole Genome Bisulfite Sequencing by Zymo**

Samples were processed and analyzed using the Methyl-MaxiSeq®: whole genome bisulfite sequencing (Zymo Research, Irvine, CA). A quick-DNA Miniprep Plus Kit (Zymo Research, Irvine, CA) was used for DNA extraction. 100 ng of genomic DNA was bisulfite converted using EZ DNA Methylation-Lightning Kit (Cat. D5031) from Zymo Research according to the manufacture's protocol. This was followed by a second strand synthesis reaction and then addition of adapters through tagmentation using Illumina's Nextera® kit. PCR was performed with Illumina Nextera Unique Dual Indices. Library quality control was performed on the Agilent 2200 TapeStation. Libraries were sequenced on an Illumina NovaSeq 6000 instrument (150 bp PE reads).

**Initial Bioinformatics Analysis**

Sequence reads from WGBS libraries were identified using standard Illumina base calling software. For pre-processing and quality control, input sequencing reads were trimmed using Trim Galore. Specific trimming parameters (e.g., directionality) were set appropriately based on the library preparation protocol. To specify, WGSB libraries prepared at Zymo Research are non-directional. Raw FASTQ files were adapter and quality trimmed, and 15 bases were further trimmed off at the 5' end according to the Nextera recommendations. Post-trimming quality control was done using FastQC. Picard tools were used to calculate the library insert size distribution.

To process WGBS data, the trimmed reads were aligned to a specified reference genome assembly using Bismark. Methylation ratios for each cytosine in CpG context were called using MethylDackel. The methylation level of each sampled cytosine was estimated as the number of reads reporting a C, divided by the total number of reads reporting a C or T. Read depths per cytosine in the whole genome as well as in different genomic regions (e.g. gene body, promoter, CpG island, etc. based on available annotations) were calculated and tabulated using scripts.

Finally, a custom MultiQC report in an interactive HTML format was generated. This final report collects QC metrics, summary data tables, visualizations, and direct download links for the raw and processed sequencing data files.

**Differential Methylation Analysis**

A comparative statistical analysis was performed to identify, annotate, and visualize differential methylated sites and regions using DSS. The data was pre-filtered to exclude low coverage cytosines and keep cytosines with read depth ≥ 5 in ≥ 2 samples in any group. DMCs and DMRs were detected using DSS and the Wald test and the Benjamini-Hochberg P-value adjustment. Significant DMCs and DMRs have FDR ≤ 0.05 and absolute methylation difference ≥ 0.1. The DMRs were considered if they had a minimum of 4 CpG sites. Annotation of DMRs was accomplished by overlapping each DMR with other functional regions, including genes, exons, introns, promoters, and CpG islands. The functional regions were derived from the UCSC or NCBI database. The minimum size for an overlap is 1 bp. Then the program g: Profiler was used to perform a functional enrichment analysis for genes linked to the DMRs. Linked genes are the set of all genes which overlap with DMRs. Circos software was used to visualize chromosomes and DMRs (https://circos.ca/software/download/circos/).

**RNA Extraction**

RNA extraction, library preparations, and sequencing reactions were conducted at Azenta (South Plainfield, NJ, USA). Total RNA was extracted from fresh frozen cell pellets using Qiagen RNeasy Plus Universal mini kit following manufacturer's instructions (Qiagen, Hilden, Germany).

**RNA Library Preparation and Sequencing**

RNA samples were quantified using Qubit 2.0 Fluorometer (ThermoFisher Scientific, Waltham, MA, USA) and RNA integrity was checked with 4200 TapeStation (Agilent Technologies, Palo Alto, CA, USA).

rRNA depletion sequencing library was prepared by using QIAGEN FastSelect rRNA HMR Kit (Qiagen, Hilden, Germany). RNA sequencing library preparation uses NEBNext Ultra II RNA Library Preparation Kit for Illumina by following the manufacturer's recommendations (NEB, Ipswich, MA, USA). Briefly, enriched RNAs are fragmented for 15 minutes at 94 °C. First strand and second strand cDNA are subsequently synthesized. cDNA fragments are end repaired and adenylated at 3'ends, and universal adapters are ligated to cDNA fragments, followed by index addition and library enrichment with limited cycle PCR. Sequencing libraries were validated using the Agilent Tapestation 4200 (Agilent Technologies, Palo Alto, CA, USA), and quantified using Qubit 2.0 Fluorometer (ThermoFisher Scientific, Waltham, MA, USA) as well as by quantitative PCR (KAPA Biosystems, Wilmington, MA, USA).

The sequencing libraries were multiplexed and clustered on a flowcell. After clustering, the flowcell was loaded on the Illumina NovaSeq 6000 instrument according to manufacturer's instructions. The samples were sequenced using a 2x150 Pair-End (PE) configuration Raw sequence data (.bcl files) generated from Illumina NovaSeq was converted into fastq files and de-multiplexed using Illumina bcl2fastq program version 2.20. One mismatch was allowed for index sequence identification.

**Analysis**

After demultiplexing, sequence data was checked for overall quality and yield. Then, sequence reads were trimmed to remove possible adapter sequences and nucleotides with poor quality using Trimmomatic v.0.36. The trimmed reads were mapped to the reference genomes using the STAR aligner v.2.5.2b. The STAR aligner is a splice aware aligner that detects splice junctions and incorporates them to help align the entire read sequences. BAM files were generated as a result of this step. Unique gene hit counts were calculated by using featureCounts from the Subread package v.1.5.2 Only unique reads within exon regions were counted.

After extraction of gene hit counts, the gene hit counts table was used for downstream differential expression analysis. Using DESeq2, a comparison of gene expression between the customer-defined groups of samples was performed. The Wald test was used to generate p-values and log2 fold changes. Genes with p-value < 0.05 were called as differentially expressed genes for each comparison. We did not apply multiple hypothesis testing correction to the RNA-seq data. As a result, the reported p-values are unadjusted and may include false positives due to the high number of comparisons. This approach was taken to allow exploratory identification of potential targets. Heatmaps for gene expression were created using the ggplot2 R-packages (4.4.1 version). Bar graphs and volcano plots were created using GraphPad Prism 9 software.

**Methyl-seq and RNA-seq data integration**

Genes with differential methylation status in their promoters or gene bodies identified in the Methyl-seq differential analysis were cross-examined with their gene expression profiling. The cut-off for differentially expressed genes was p-value <0.1.

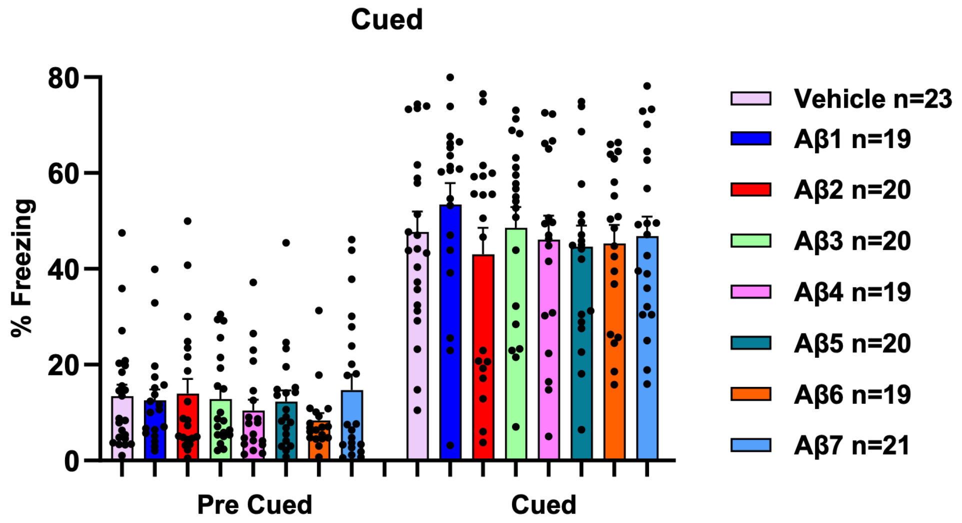


**Fig. S1: Prolonged exposure to pM Aβ does not affect cued fear conditioning.** Cued fear memory does not show differences (Cued: 1-way ANOVA F (7,153) = 0.4708, p = 0.8546; Pre Cued: 1-way ANOVA F (7,153) = 0.6544, p = 0.7102, Bonferroni's $p > 0.9999$ for every condition).

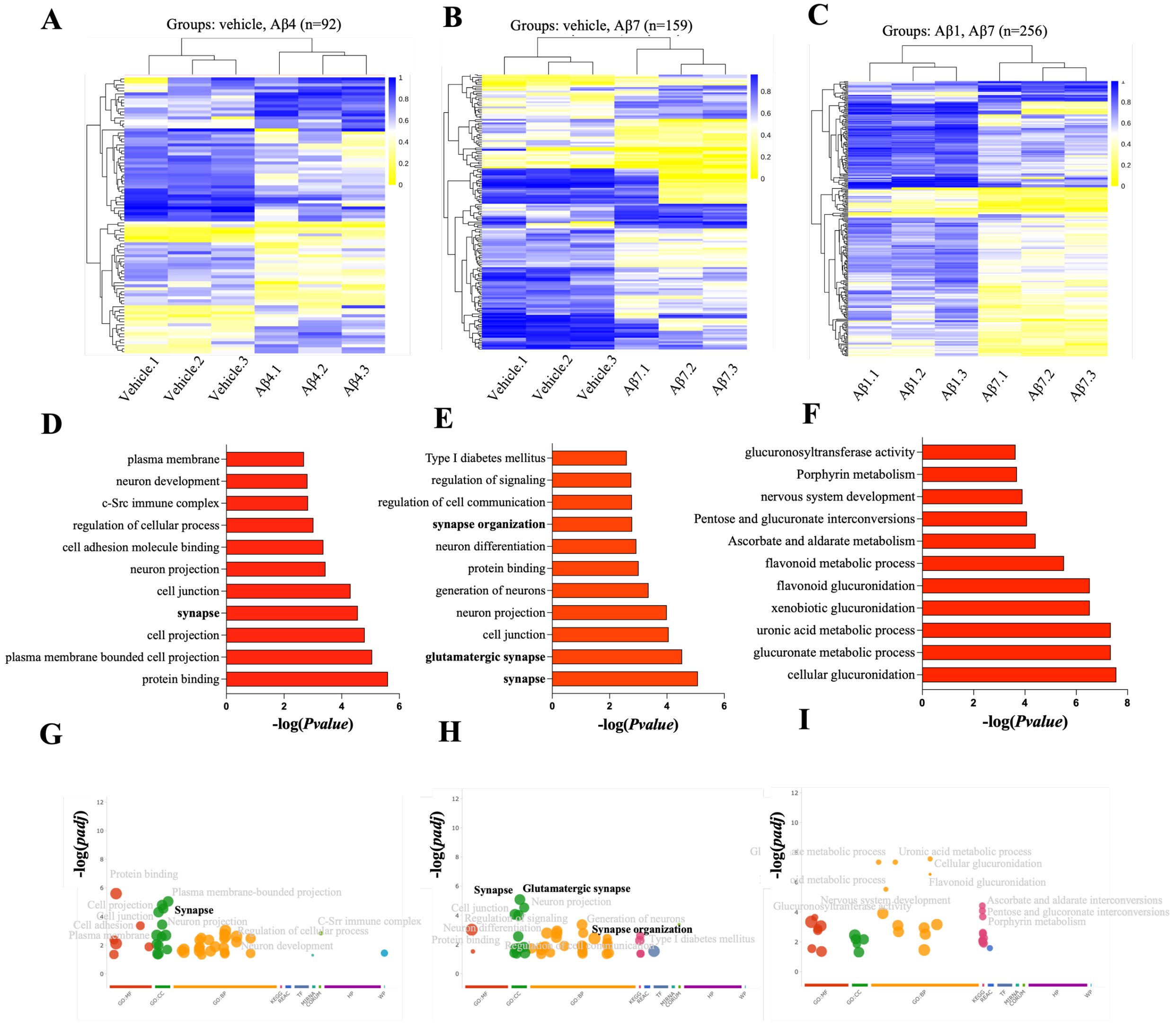


**Fig. S2: Picomolar Aβ affects DNA methylation differently at distinct time points.** (A-C) Cluster heatmaps of DMRs for Vehicle/Aβ4 (A), Vehicle/Aβ7 (B), Aβ1/Aβ7 (C) (FDR ≤ 0.05 and absolute methylation difference ≥ 0.1). (D-F) GO enrichment analysis for Vehicle/Aβ4 (D), Vehicle/Aβ7 (E), Aβ1/Aβ7 (F). Pathways are shown in ascending order based on $-\log_{10}$ (Pvalue). (J-L) Manhattan plots for Vehicle/Aβ4 (G), Vehicle/Aβ7 (H), Aβ1/Aβ7 (I). The x-axis represents functional terms that are grouped and color-coded by data sources (e.g. Molecular Function from GO is red). The y-axis shows the adjusted enrichment p-values in negative log10 scale. GO terms related to the synapse are highlighted in bold letters.

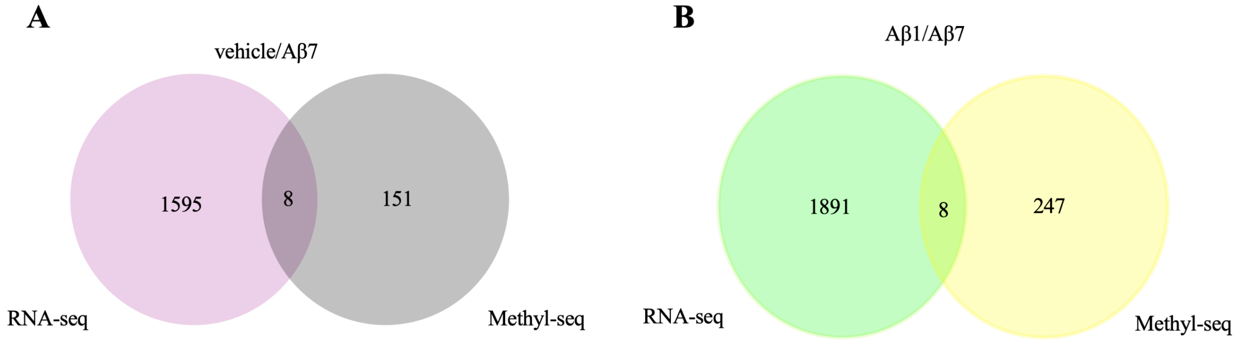


**Fig. S3: Integrated analysis of DMRs and DEGs following exposure to pM Aβ for vehicle/Aβ7, Aβ1/Aβ7**. (A-B) Venn diagrams representing the overlap of DEGs and DMRs in vehicle/Aβ7 (A), Aβ1/Aβ7 (B). $p \leq 0.1$.

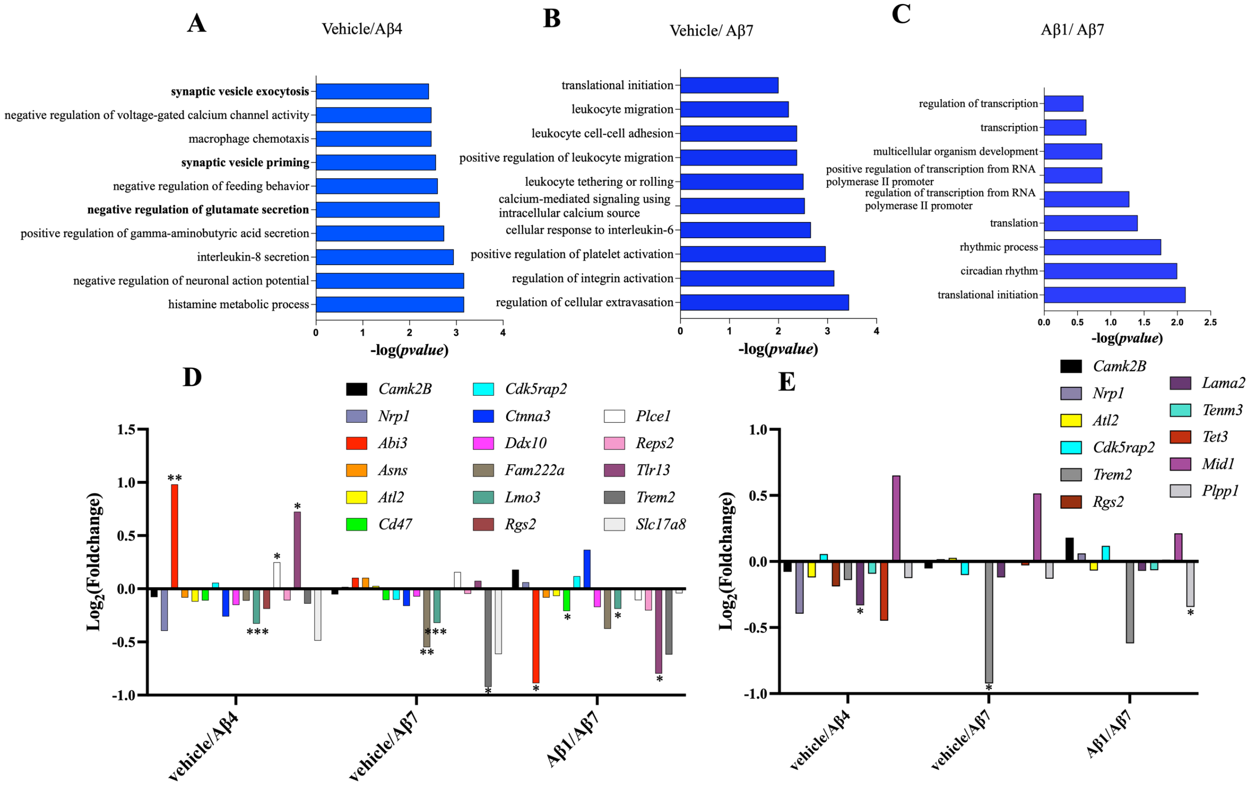


**Fig. S4: Changes in RNA expression in genes that are differentially methylated following exposure to pM Aβ for Vehicle/Aβ4, Vehicle/Aβ7, Aβ1/Aβ7. (A-C)** GO enrichment analysis for Vehicle/Aβ4 (A), Vehicle/Aβ7 (B), Aβ1/Aβ7 (C). Pathways are shown in ascending order based on $-\log_{10}$ (Pvalue). (D-E) Histogram representing the DEGs associated with AD (D) and synaptic function (E) found differentially methylated and expressed in at least one condition. *, $p \leq 0.05$, **, $p \leq 0.01$; ***, $p \leq 0.001$.

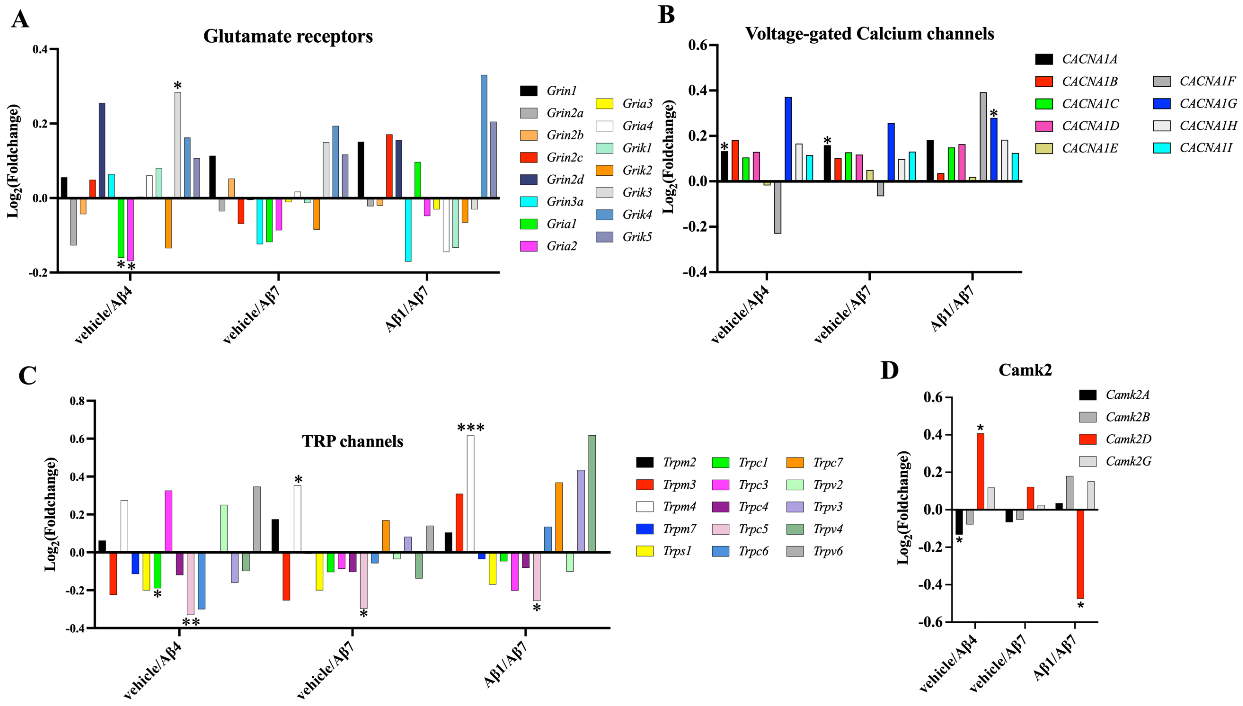


**Fig. S5. Changes in RNA expression in genes involved in $Ca^{2+}$ homeostasis following exposure to pM Aβ for Vehicle/Aβ4, Vehicle/Aβ7, Aβ1/Aβ7. (A-D)** Histogram representing the genes encoding glutamate receptors (A), Voltage-gated calcium channels (B), TRP channels (C), and Camk2 (D). *, $p \leq 0.05$, **, $p \leq 0.01$, ***, $p \leq 0.001$.

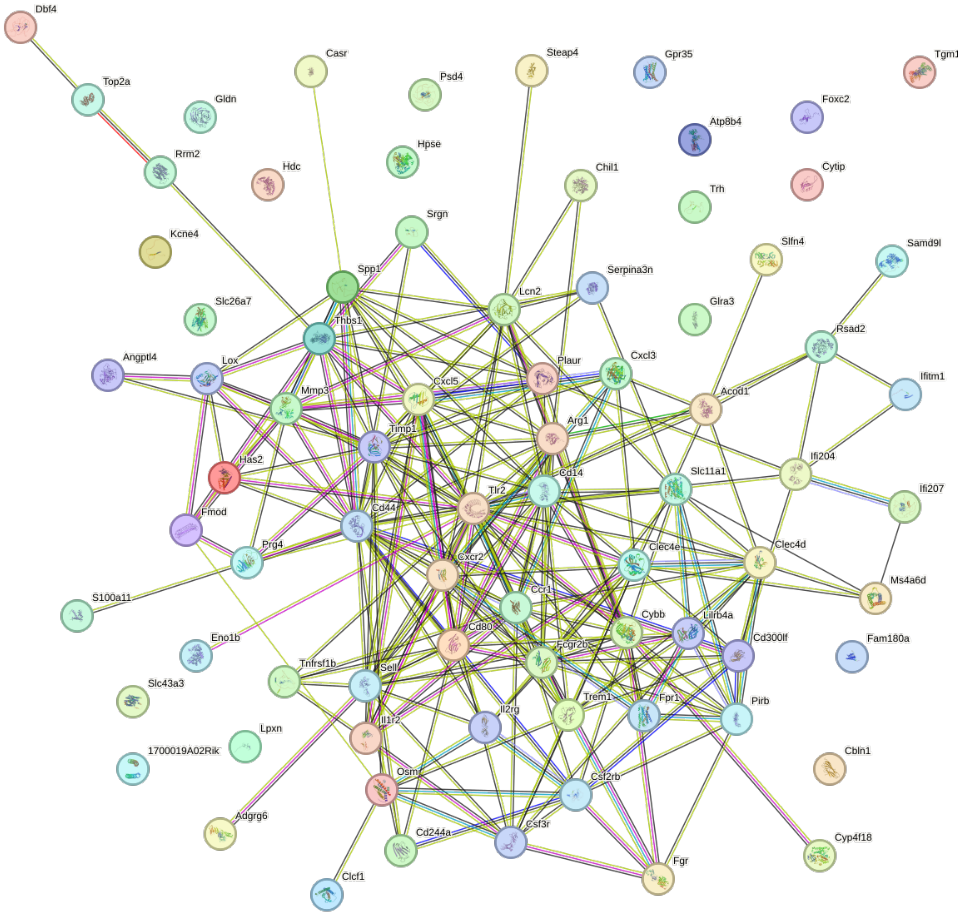


**Fig. S6: Pathways analysis of DEGs upregulated following exposure to pM Aβ for vehicle/ Aβ1.** STRING analysis was performed on the genes found differentially expressed with $log_2$(fold change) ⩾ 1 and p ⩽ 0.05. For the analysis, text mining, experiments, and databases were chosen for active interaction sources. A value of 0.400 was selected as the minimum required interaction score. For the analysis, text mining, experiments, and databases were chosen for active interaction sources, and a value of 0.400 was selected as the minimum required interaction score. Line colors represent known interactions from curated databases (blue), experimentation (purple), gene neighborhood (green), gene fusions (red), gene co-occurrence (blue), text mining (yellow), and co-expression (black).

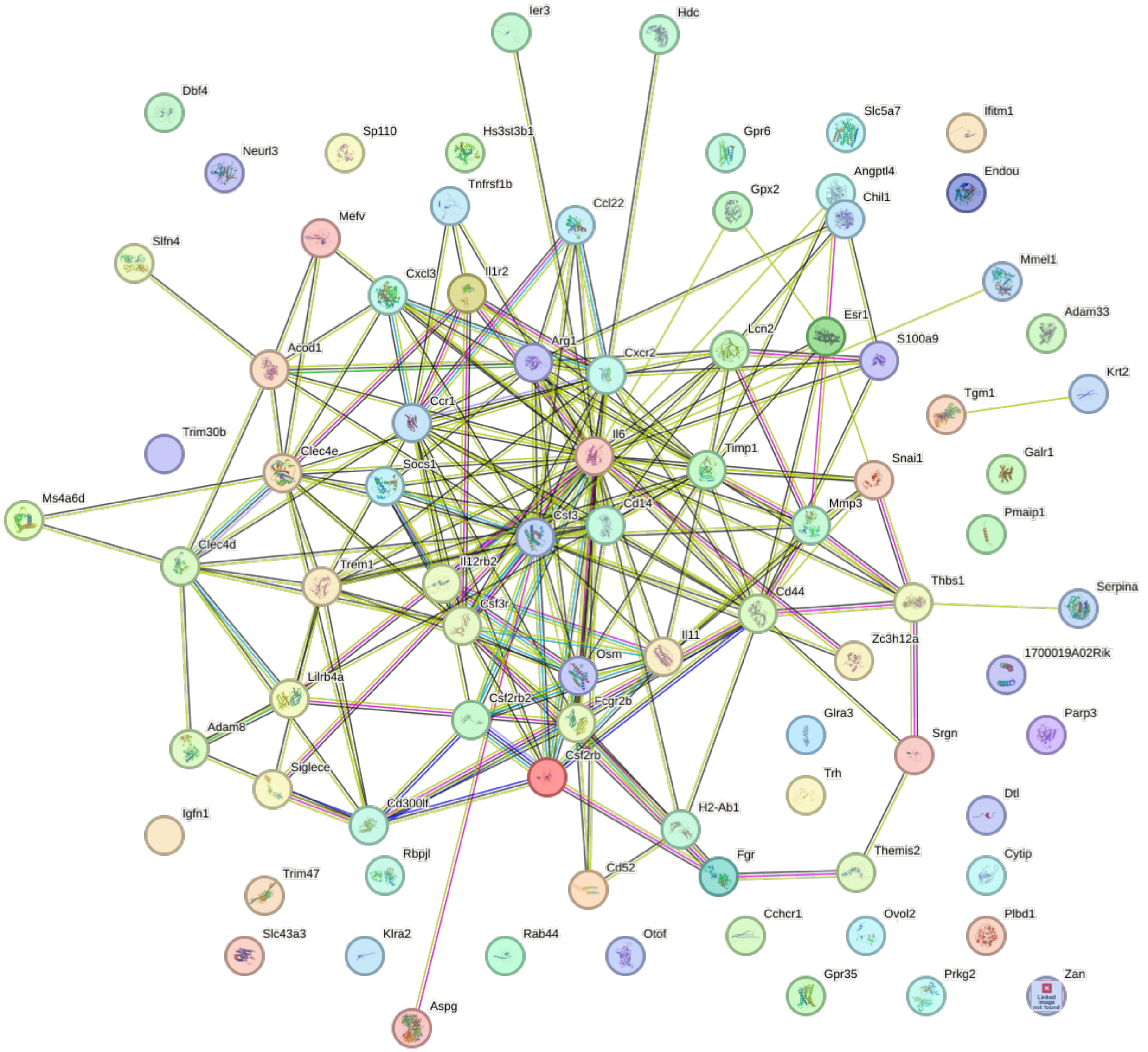


**Fig. S7: Pathways analysis of DEGs upregulated following exposure to pM Aβ for vehicle/ Aβ4.** STRING analysis was performed on the genes found differentially expressed with $\log_2$(fold change)$\geq$ 1 and p$\leq$0.05. For the analysis, text mining, experiments, and databases were chosen for active interaction sources. A value of 0.400 was selected as the minimum required interaction score. For the analysis, text mining, experiments, and databases were chosen for active interaction sources, and a value of 0.400 was selected as the minimum required interaction score. Line colors represent known interactions from curated databases (blue), experimentation (purple), gene neighborhood (green), gene fusions (red), gene co-occurrence (blue), text mining (yellow), and co-expression (black).

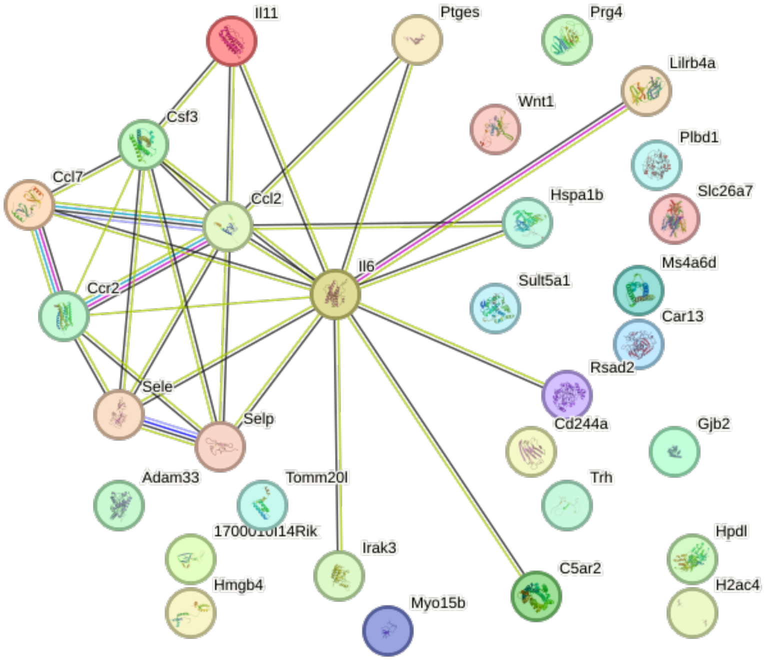


**Fig. S8: Pathways analysis of DEGs upregulated following exposure to pM Aβ for vehicle/ Aβ7.** STRING analysis was performed on the genes found differentially expressed with $log_2$(fold change)≥1 and p≤0.05. For the analysis, text mining, experiments, and databases were chosen for active interaction sources. A value of 0.400 was selected as the minimum required interaction score. For the analysis, text mining, experiments, and databases were chosen for active interaction sources, and a value of 0.400 was selected as the minimum required interaction score. Line colors represent known interactions from curated databases (blue), experimentation (purple), gene neighborhood (green), gene fusions (red), gene co-occurrence (blue), text mining (yellow), and co-expression (black).

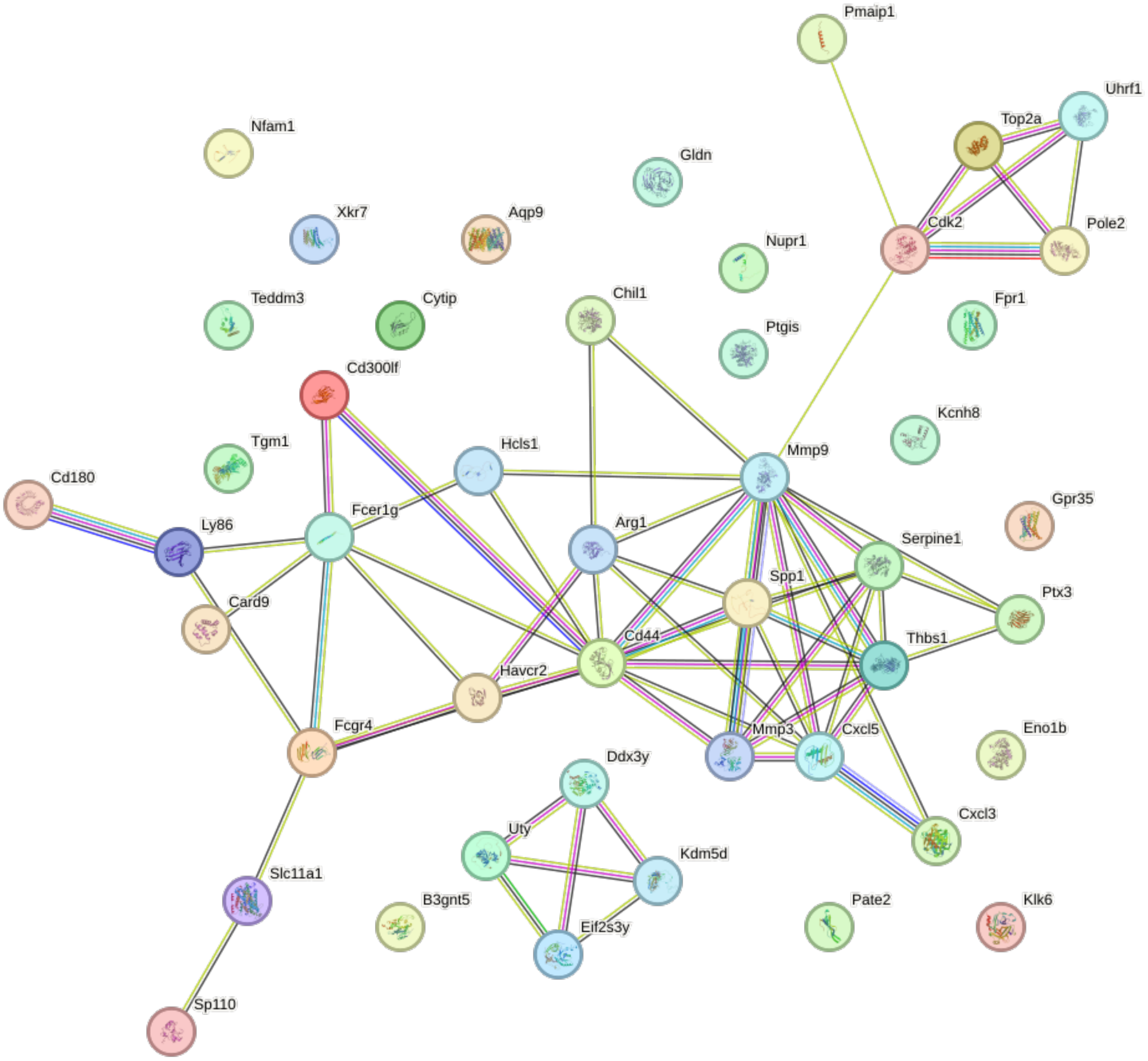


**Fig. S9: Pathways analysis of DEGs downregulated following exposure to pM Aβ for Aβ1/Aβ7.** STRING analysis was performed on the genes found differentially expressed with $\log_2$(fold change) ≤1 and p ≤ 0.05. For the analysis, text mining, experiments, and databases were chosen for active interaction sources. A value of 0.400 was selected as the minimum required interaction score. For the analysis, text mining, experiments, and databases were chosen for active interaction sources, and a value of 0.400 was selected as the minimum required interaction score. Line colors represent known interactions from curated databases (blue), experimentation (purple), gene neighborhood (green), gene fusions (red), gene co-occurrence (blue), text mining (yellow), and co-expression (black).

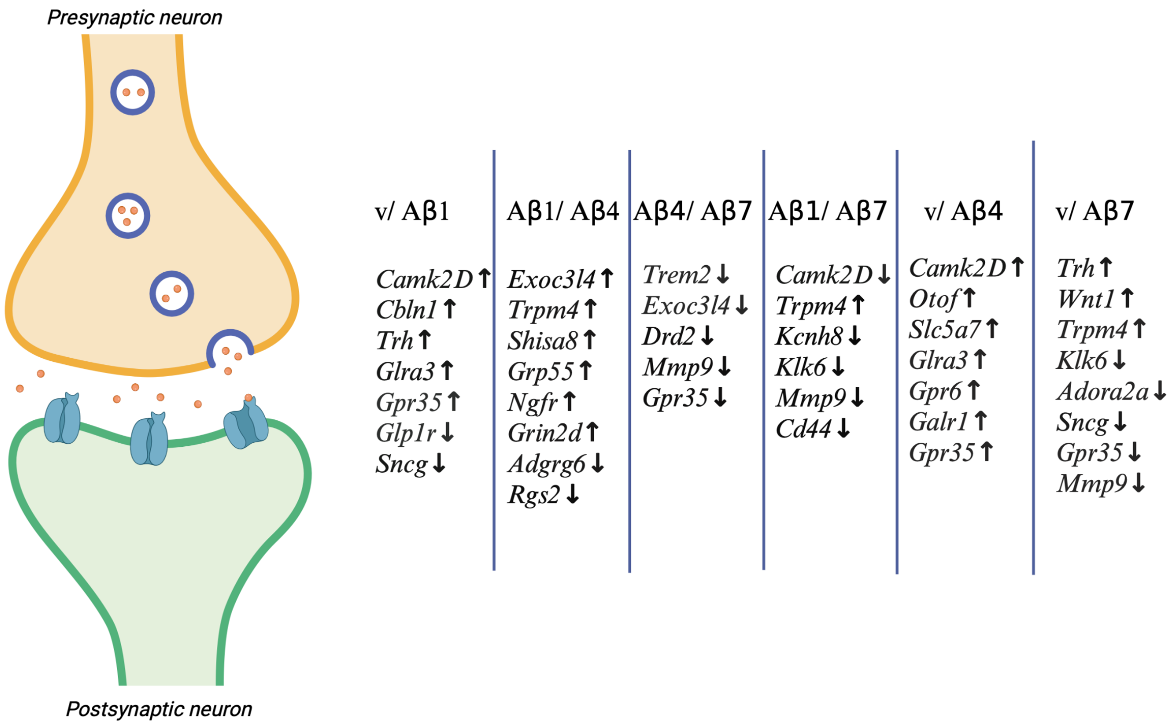


**Fig. S10: Illustration of DEGs following pM Aβ exposure associated with synaptic function and $Ca^{2+}$ homeostasis.** Illustration showing the DEGs involved in pre- or postsynaptic function in vehicle/Aβ1, Aβ1/Aβ4 and Aβ4/Aβ7, Aβ1/Aβ7, vehicle/Aβ4 and vehicle/Aβ7 having $p \leq 0.05$ and $|\log2foldchange| \geq 1$ and DEGs involved in $Ca^{2+}$ homeostasis having $p \leq 0.05$ and $|\log2foldchange| \geq 0.4$. Created with BioRender.com released under a Creative Commons Attribution-NonCommercial-NoDerivs license.
